# Evolutionary conservation of maternal RNA localization in fishes and amphibians revealed by TOMO-Seq

**DOI:** 10.1101/2021.08.01.454628

**Authors:** Ravindra Naraine, Viktoriia Iegorova, Pavel Abaffy, Roman Franek, Vladimír Soukup, Martin Psenicka, Radek Sindelka

## Abstract

Asymmetrical localization of biomolecules inside the egg, results in uneven cell division and establishment of many biological processes, cell types and the body plan. However, our knowledge about evolutionary conservation of localized mRNAs is still limited to a few candidates. Our goal was to compare localization profiles along the animal-vegetal axis of mature eggs from four models, *Xenopus laevis*, *Danio rerio*, *Ambystoma mexicanum* and *Acipenser ruthenus* using the spatial expression method called TOMO-Seq. We revealed that RNAs of many known important genes such as germ layer determinants, germ plasm factors and members of key signalling pathways, are localized in completely different profiles among the models. There was also a poor correlation between vegetally localized genes but a relatively good correlation between animally localized genes. These findings indicate that the regulation of embryonic development within the animal kingdom is highly diverse and cannot be deduced based on a single model.

## Introduction

One of the most intriguing question in biology is how the existence of asymmetrical biomolecule localization in the oocytes and early embryos influences the developmental process. Several recent studies have utilized RNA-Seq to completely characterize the spatial transcriptome using eggs and embryos from popular model organisms such as the *Xenopus laevis* (Owens et al. 2017; Sindelka et al. 2018) and *Danio rerio* (Junker et al. 2014). However, a comprehensive comparative study of the evolutionary development between species among different groups is still missing.

During evolution, the first sign of separation from fishes with fins to tetrapodomorpha happened approximately 380 million year ago, during the Devonian period (or Age of Fishes). At around 350 million years ago, the first walking tetrapods evolved from their aquatic ancestors. Animals that belong to bony vertebrates fall into two groups: the *Actinopterygii* (ray-finned fishes) and the *Sarcopterygii* (lobe-fined fishes) which includes the *Tetrapoda* subgroups (amphibians, reptiles, birds and mammals). The sterlet (*Acipenser ruthenus*), a member within the Chondrostei branch, and the Teleostei fish - zebrafish (*D. rerio*), both belong to the *Actinopterygii* group. The tetrapods (amphibians such as *X. laevis* and *Ambystoma mexicanum*) and the amniotes have both evolved from the *Sarcopterygii* branch (Fig. 1a) (Wourms 1997; Volff 2005; Clack 2012; Yamamoto et al. 2017; Wake and Koo 2018).

**Figure 1.**
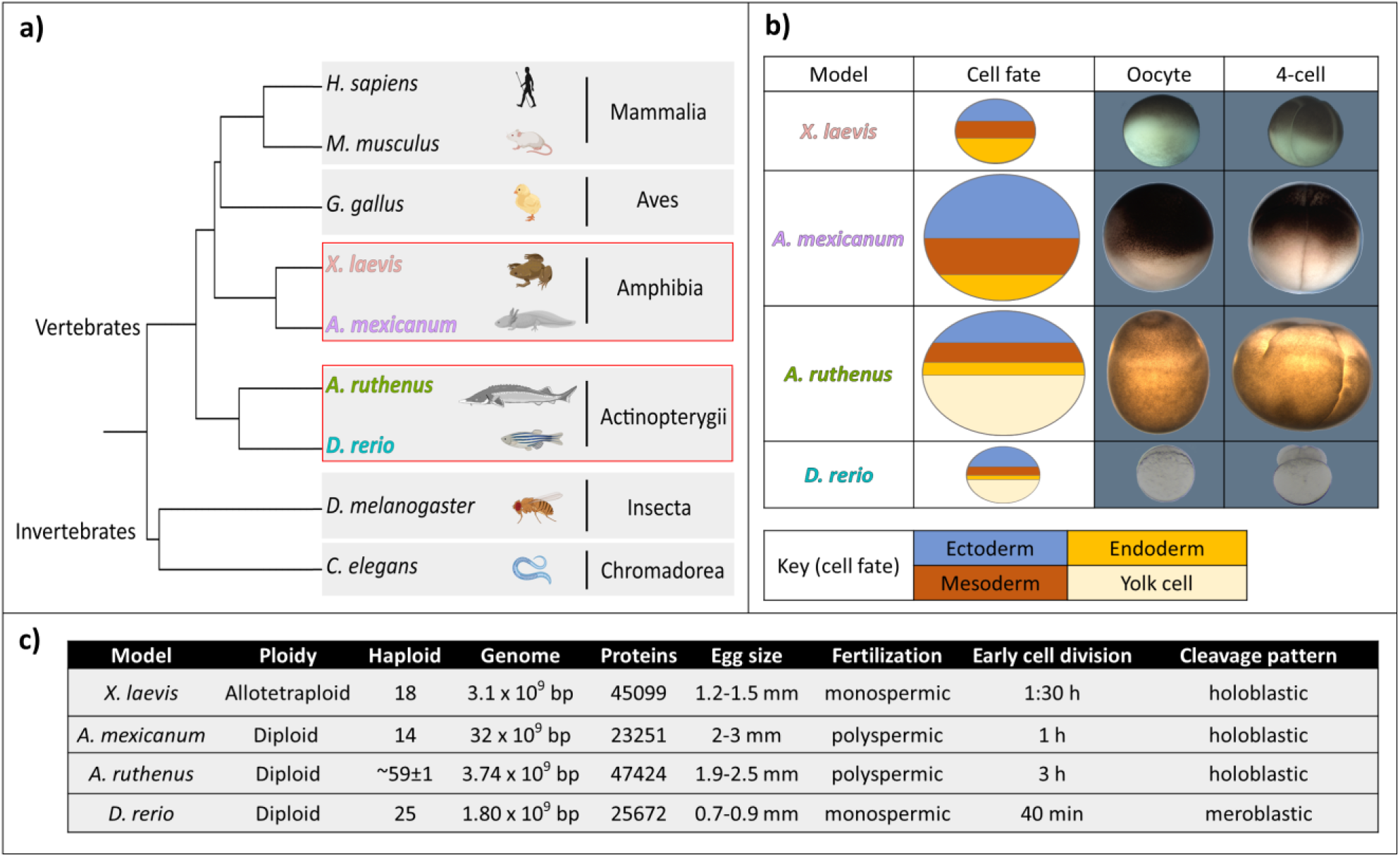
Relationship between the studied taxa and their developmental features. a) Taxonomic tree based on NCBI taxonomic lineage of select animal models. Highlighted are the model organisms used in this research. b) Images of the oocyte, 4-cell stages and cell fate maps for the analysed models. Images have been cropped and brightness adjusted for clarity. c) Genomic and egg developmental properties for each model.

Amphibians and ray-finned fishes show great variations in their body shapes, sizes and also in the mechanisms driving their early development, such as germ-layer determination, yolk storage and germ-cell specification. *Acipenseridae* are usually long-lived fishes with slow growth and maturation (Pikitch et al. 2005). Representatives of this family grow continuously with age, with some of them such as the *A. ruthenus* becoming matured within 3-9 years (Chebanov and Galich 2013), and can live up to 24 years with a maximum size of 125 cm and weight up to 16 kg (FAO Fisheries & Aquaculture). Contrasting to *A. ruthenus*, the *D. rerio* is a smaller fish that grows to a maximum size of 3.5 cm and reaches maturation within a short period of 2.5 months (Eaton and Farley 1974; Spence et al. 2007). Representatives of the *Amphibia* class: African-clawed frog (*X. laevis*) matures within 1-2 years with the adult reaching sizes of up to 10 cm (Xenbase) and axolotl (*A. mexicanum*) which is locked at the larval stage, reaches the sexual maturity at approx. 12 months, and can grow up to 30 cm (Gresens 2004). There is also variation amongst the sizes (diameter) of the matured eggs (*D. rerio*: 0.7-0.9 mm, *X. laevis*: 1.2-1.5 mm, *A. ruthenus:* 1.9-2.5 mm, *A. mexicanum*: 2-3 mm) (Fig. 1b) (Dettlaff et al. 1981; Dettlaff and Rudneva 1991; Kimmel et al. 1995; Gresens 2004; Uusi-Heikkilä et al. 2010; Chebanov and Galich 2013) and also in the fertilization method. In *D. rerio*, monospermic fertilization is observed, whereby only one spermatozoa penetrates into the egg through the only one existing micropyle in the animal pole (Joo and Kim 2013). In contrast, fertilization in *A. ruthenus* is characteristically physiological polyspermy (penetration of numerous spermatozoa into an egg through numerous micropyles) (Zalenskii 1878; Iegorova et al. 2018). Eggs from *X. laevis* undergo typical monospermic fertilization, while those from *A. mexicanum* display polyspermic fertilization (Bordzilovskaya and Dettlaff 1991; Dettlaff and Rudneva 1991).

Genome sizes of the above listed animals are different too (Fig. 1c): *D. rerio* - 1.8 Gb (Hinegardner and Rosen 1972), *X. laevis* - 3.1 Gb (Hirsch et al. 2002) and *A. ruthenus* - 3.74 Gb (Birstein et al. 1993). Interestingly, the genome size of *A. mexicanum* is approximately 32 Gb (Keinath et al. 2015), which is caused by the expansion of introns and intergenic regions. Ploidies are varying too: *A. mexicanum*, *A. ruthenus* and *D. rerio* are diploids, while *X. laevis* is allotetraploid with a haploid numbers: 14, 59±1, 25 and 18, respectively (Frankhauser and Humphrey 1942; Post 1965; Birstein and Vasiliev 1987; Ludwig et al. 2001; Hirsch et al. 2002; Menon et al. 2017). Differences can also be observed in the division time during early embryogenesis, from tens of minutes for *D. rerio*, while hours for *X. laevis*, *A. mexicanum* and *A. ruthenus* (Xenbase; Dettlaff et al. 1981; Bordzilovskaya and Dettlaff 1991; Kimmel et al. 1995). Most of the vertebrate eggs share animal–vegetal axis polarization (King et al. 2005; Houston 2017). However, during early development the cleavage patterns vary. *Acipenser ruthenus* and amphibian embryos undergo holoblastic cleavage (completely dividing embryo), while teleosts (including *D. rerio*) exhibit a meroblastic pattern (most of the yolk mass remains uncleaved at the blastula stage) (Fig. 1b) (Ballard 1981; Bordzilovskaya and Dettlaff 1991; Kimmel et al. 1995; Elinson 2009). *Acipenseridae* and *Amphibia* also share strong cytological similarities during their oogenesis, such as: nucleolar and other nuclear structures, cytoplasmic organelles, the same structure of yolk platelets, presence of cortical granules, absence of ribosomes in previtellogenesis, extrusions of nucleolar material into the cytoplasm, and the same dense material cementing the mitochondria (Raikova 1973, 1974).

Similarities and differences between the eggs and embryogenesis are propagated in the particular distribution of biomolecules regulating development (King et al. 2005; Kloc and Etkin 2005). Proteins, RNAs, cellular structures and organelles are distributed unevenly within the egg (Marlow 2010). During early oogenesis, the oocytes of many animals contain an organelle called the Balbiani body (or mitochondrial cloud), which plays an important role in the transportation of the germ cell determinants and the distribution of RNAs to the oocyte’s vegetal cortex (Kloc et al. 2004; Marlow 2010). RNA localization in *X. laevis* also controls embryonic patterning and determines the specification along the animal-vegetal (A-V) axes, which later defines where the three primary germ layers will originate (Flachsova et al. 2013; Owens et al. 2017). These three germ layers consist of the: endoderm (gastrointestinal, respiratory and urinary systems) which develops from the vegetal part, mesoderm (notochord, axial skeleton, cartilage, connective tissue, trunk muscles, kidneys and blood) which develops in the meridial part, and ectoderm (nervous system, epidermis and various neural crest-derived tissues) which develops from the animal part in *X. laevis* (Kiecker et al. 2016). A similar early development can also be observed in *A. mexicanum* (Pasteels 1942). Contrastingly the endoderm of the *A. ruthenus* develops in the meridian area while the meso- and ectoderm layers shift towards the animal part compared to *X. laevis*. The vegetal part of *A. ruthenus* contains the primordial germ cell factors and also the yolk, which is used as an extra-embryonic tissue with nutritional function (Dettlaff et al. 1993; Saito et al. 2014; Pocherniaieva et al. 2018). The *D. rerio* fate map closely resembles that of the *A. ruthenus* (Fig. 1b), whereby the early development occurs mainly at the animal pole region, while the vegetal region is dedicated as yolk storage (Fig. 1b) (Ober et al. 2003).

Most of the studies that focused on RNA localization were performed using the *X. laevis* eggs and primarily targeted the genes within the vegetal pole. These studies found that RNA localization happens during oogenesis due to early and late transportation pathways (Kloc and Etkin 1995). mRNAs coding *nanos1* (*xcat2*) and *dazl* are distributed to the vegetal pole by the early pathway and play an important role in the determination and migration of the primordial germ cells (PGCs). Other vegetally localized mRNAs, such as *gdf1* (previously *vg1)*, *vegt* and *wnt11,* have been found to be mesodermal and endodermal determinants, and are also crucial for left-right axis formation in the embryo (Forristall et al. 1995; Gilbert 2000; King et al. 2005; Kloc and Etkin 2005). Saito et al. (2014) demonstrated that in *A. ruthenus*, PGCs are formed in the vegetal pole as well. In contrast to *X. laevis* and *A. ruthenus*, urodele (*A. mexicanum*) embryos stimulate PGC formation from the primitive ectoderm (the animal cap) by induction in the absence of germ plasm, and their formation is more similar to mammals than to anurans (Sutasurja and Nieuwkoop 1974; Bachvarova et al. 2004). Theusch et al. (2006) described a different mechanism in the early embryos of the *D. rerio*, where *vasa*, *nanos1* and *dnd1* mRNAs, form part of the germ plasm and are present in a wide cortical band at the animal pole and belong to the first class of germ RNA components. *Dazl* mRNAs also participate in germ plasm formation. However, they belong to the second RNA class, which is localized on the vegetal cortex of the freshly laid egg and then later migrates towards the animal pole after fertilization. These RNAs exhibit separate pathways of segregation, which leads to a characteristic substructure of the germ plasm.

In this study, we utilized spatial RNA sequencing (TOMO-Seq) to determine RNA localization along the animal-vegetal axis of fully grown eggs from two representatives of *Actinopterygii* (*A. ruthenus*, *D. rerio*) and two representatives of *Amphibia* (*X. laevis* and *A. mexicanum*) species. We focused especially on genes (to simplify, human orthologs were identified and human symbols are used in remaining text) with known functions that are part of the germ cell, germ layer determinants and developmentally important signalling pathways. We determined their evolutionary conservation and analysed for the presence of putative localization motifs.

## Results

### Global asymmetric mRNA localization is a general feature of fish and amphibian eggs

TOMO-Seq analysis was performed using mature eggs from four animal models (*X. laevis*, *A. mexicanum*, *A. ruthenus* and *D. rerio*), which were dissected into five constitutive segments along their animal-vegetal axis (A- extremely animal, B- animal, C- central, D- vegetal, E- extremely vegetal). The relative proportion of the concentration of the RNA extracted from each section for each model is shown in the Supplemental file 1: Fig. S1. Sequences were annotated based on the known reference transcriptomes (*X. laevis*, *A. mexicanum* and *D. rerio*) or against the *de novo* transcriptome (*A. ruthenus*). When possible, the mRNA sequences were given a gene nomenclature based on the human gene symbols of its most probable human ortholog match.

We created three datasets from the analyzed data (Supplemental file 2: Table S1). Dataset1 comprised of all the DEGs and was used for a less stringent ortholog and localization comparative analysis between models, paralog analysis within the same model and comparative GO analysis. Dataset2 comprised of reproducible DEGs with well-defined profiles and was used for motif enrichment analysis and detection of known motifs. Dataset3 comprised of a smaller subset of curated DEGs with well defined, reproducible profiles followed by additional annotation analysis, and was used to carry out a more comprehensive ortholog comparative analysis between the models.

We identified in total, the following number of maternal genes in *X. laevis*: 27889, *A. mexicanum*: 32960, *A. ruthenus*: 42303 and *D. rerio*: 15867, with an average expression higher than zero within the egg. The majority of these mRNAs for most of the models (*X. laevis*: 75%, *A. mexicanum*: 90%, *A. ruthenus*: 51% and *D. rerio*: 11%) were DEGs (Fig. 2b, Supplemental file 2: Table S1-dataset1). Each model had between 8000-11000 DEGs with a unique gene symbol (annotated) (*X. laevis*: 10753, *A. mexicanum*: 8822, *A. ruthenus*: 9590), except for *D. rerio* (1422). Of these annotated DEGs, 980 were shared amongst all models, 7578 between the amphibians and 1223 between the fishes (Supplemental file 1: Fig. S2). Each model, except for *D. rerio*, shared a similar pairwise comparative number (between 6000-8000) of annotatable DEGs, which equates to between 71-86% of the organism’s annotatable maternal DEGs. Despite only 11-14% of each model’s annotatable DEGs matching those from *D. rerio*, they represented a large portion 82-89% of the identified *D. rerio* annotatable maternal DEGs.

**Figure 2.**
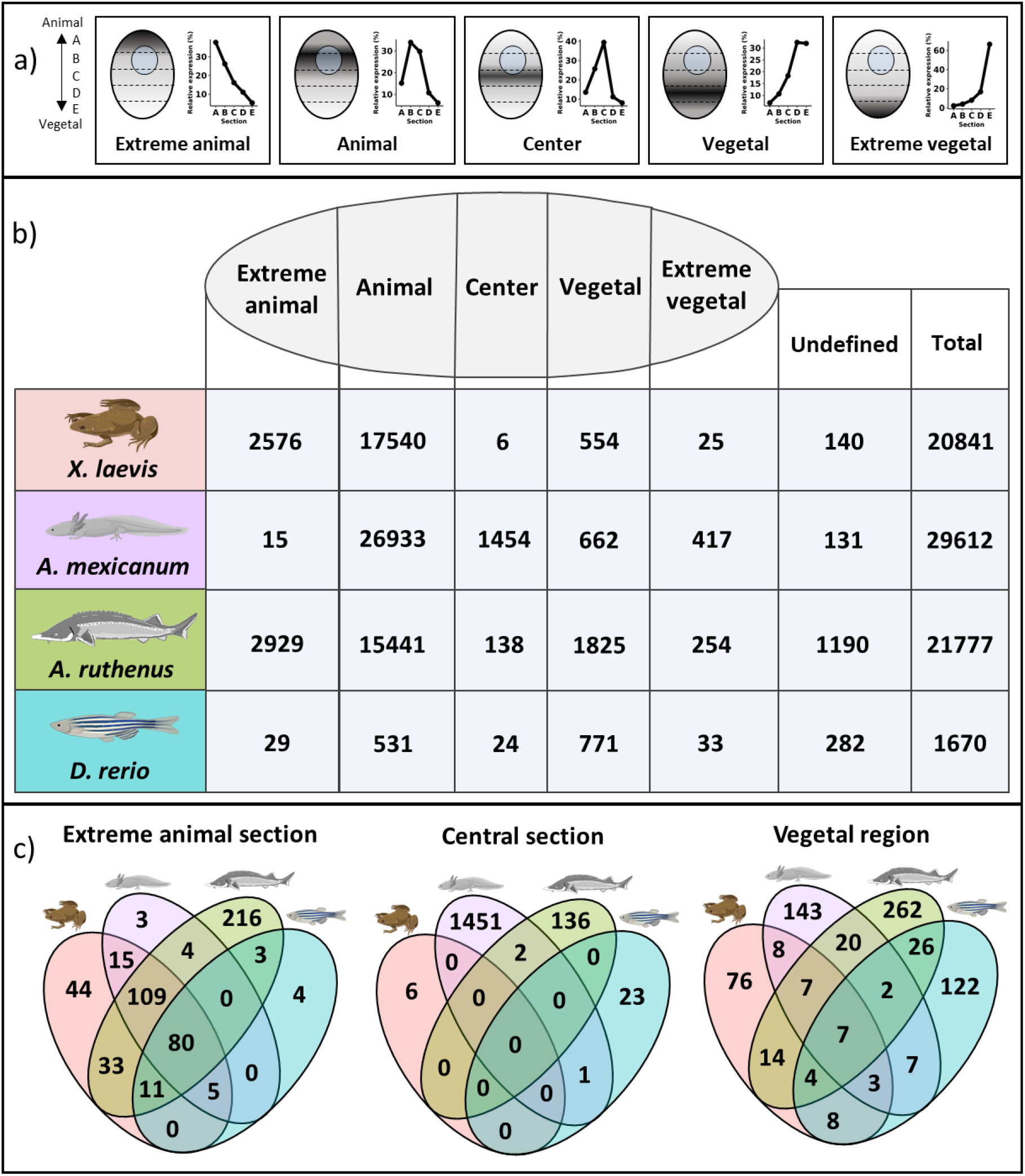
Localization of maternal differentially expressed genes. a) Schematic of the five main localization profiles detected for the maternal genes within the egg. The darker colored regions within the egg represent higher gene count saturation. The line graphs represent the median expression of the genes within all models that showed the strongest distributions for each profile. b) Number of differentially expressed genes within each localization profiles for each model (dataset1). c) Overlap of some similar/orthologous DEGs (dataset3) that share the same localization profile. The vegetal region comprises of representatives from both the vegetal and extreme vegetal localization profiles. The extreme animal section comprises some additional animal genes from *A. mexicanum* and *D. rerio*, and also consisted of any matching orthogous animal genes. Multiple paralogous genes in one model were found to match singular orthologous genes in another model. In such a case the gene count for the matching ortholog is equated to the number of the paralogs in the other model so as to adequately represent the overlap within the Venn diagram.

Of the maternal DEGs, *X. laevis*: 99.3%, *A. mexicanum*: 99.6%, *A. ruthenus*: 94.5% and *D. rerio*: 83.1% could be classified into one of the five categories (extreme animal, animal, central, vegetal, extreme vegetal) (Fig. 2a). All of the localization profiles were identifiable within each model, but at different frequencies. A large proportion (87-99%) of the unique annotatable DEGs from each model, came from the DEGs that were localized in the extreme animal and animal regions, except for *D. rerio* where only 35% originated. Of these annotated DEGs, 64% to 86% were shared between models, except *D. rerio* where 4-5% were shared. The extreme animal profile was represented by DEGs mainly in *X. laevis*: 12% and *A. ruthenus*: 13%, while in lower proportions in *D. rerio*: 2% and *A. mexicanum*: 0.1%. The *A. ruthenus* not only had the greatest number of extreme animal DEGs, but its representatives showed the steepest profiles with the highest gene abundance being present within the section animal cap (A) versus the other sections. The majority of the DEGs could be found within the animal profile (peak in the first third from animal pole – segment B) with *X. laevis*: 84%, *A. mexicanum*: 91% and *A. ruthenus*: 71%. *Danio rerio* with 32% was an exception.

Only a low proportion of the annotatable DEGs contributed to the central genes (maximum in the middle - segment C and decreased amounts in the poles), with 0-2% for all model except *A. mexicanum* with 7%. Amongst the DEGs, *A. mexicanum* had the most at 5%, while *A. ruthenus*: 1%, *D. rerio*: 1% and *X. laevis*: 0.03%. Almost no annotatable central genes were shared between the different models.

The extreme vegetal and vegetal genes comprised of only 2-7% of the annotatable maternal DEGs for all models, except for *D. rerio* with a large 48%. About 3-17% of these genes were shared between each model. The vegetal profile, contained on average the 3^rd^ most abundance of DEGs after the animal and extreme animal profiles, except for *D. rerio* where the majority of DEGs could be found. The vegetal profile proportion for each model was as follows, *X. laevis*: 3%, *A. mexicanum*: 2%, *A. ruthenus*: 8% and *D. rerio*: 46%. The extreme vegetal profile (section E > ∼50% of mRNA in egg), similarly to the center gradient, contained very few DEGs with a distribution of *X. laevis*: 0.1%, *A. mexicanum*: 1%, *A. ruthenus*: 1% and *D. rerio*: 2%. There were no genes from the subset of genes with non-DEG status that were identifiable as being ubiquitously (mRNA between 20-30% in each egg section) localized throughout the egg amongst any of the models.

Some of the selected models are known to have evolved from ancestors that have undergone complete or partial genome duplication events. As a result, we have used the available sequence similarities or gene (model specific/human) symbols to identify the most probable duplicated genes (homologs) and their localization profiles. We identified the following number of maternal gene symbols (DEG/non-DEG) as potentially duplicated, *X. laevis*: 8972, *A. mexicanum*: 4481, *A. ruthenus*: 3332 and *D. rerio*: 95. Among them only DEGs, *X. laevis*: 438, *A. mexicanum*: 67, *A. ruthenus*: 94 and *D. rerio*: 1, showed contrasting profiles (one form has extreme vegetal/vegetal profile and the duplicated form has extreme animal/animal profile) (Supplemental file 2: Table S2). The number of duplicated forms in *A. mexicanum* is probably an order higher, however given the complexity of the data due to the presence of several unknown lowly expressed variants, only genes with a raw (before normalization) gene count greater than 30 copies in any sample were analyzed.

### Conservation of vegetally localized mRNAs is poor and species dependent

After manually filtering for the most reproducible genes showing distinct profiles, a total of *X. laevis*: 127, *A. mexicanum*: 197, *A. ruthenus*: 342 and *D. rerio*: 179 genes with vegetal and extremely vegetal mRNA localization were analyzed (dataset3) (Supplemental file 2: Table S1). The first objective looked for the level of conservation of these vegetal genes amongst the four species based on its annotatable gene symbols (Fig. 2c; Supplemental file 2: Table S1, S3). We identified seven ortholog genes with vegetal and in most cases extremely vegetal localization profiles. We also determined amphibian and fish specific vegetal genes (vegetal or extremely vegetal vs other localization). Amphibians contained eight genes responsible for negative regulation of the Wnt pathway. We also identified 26 fish specific vegetal genes. Interestingly, 16 genes were found that showed vegetal profiles in only three species while showing a completely different profile in the remaining fourth. We validated the reproducibility of this result using RT-qPCR on independently prepared samples and found that the tested genes were not vegetal or had variable results (Supplemental file 1: Fig. S3). In addition, we found that there was an even higher proportion of vegetal genes which are just species specific (*X. laevis* - 76, *A. mexicanum* - 143, *A. ruthenus* - 262 and *D. rerio* - 122) (Fig. 2c). It is clear from GO enrichment analysis that species specific genes are important for protein localization, regulation of development and many other key processes (Fig. 3). The complete list of enriched GO terms for the given categories is available in Supplemental file 2: Table S4. Analysis of the enriched GO terms, metabolic pathways, transcription factors and protein complexes for all extreme vegetal/vegetal DEGs, showed a large overlap (358) of similar terms between the models. The most interesting of these, appear to be genes involved in protein binding, the epididymis, endometrium, fallopian, testis, and transcription factors involved in differentiation and cell cycle control.

**Figure 3.**
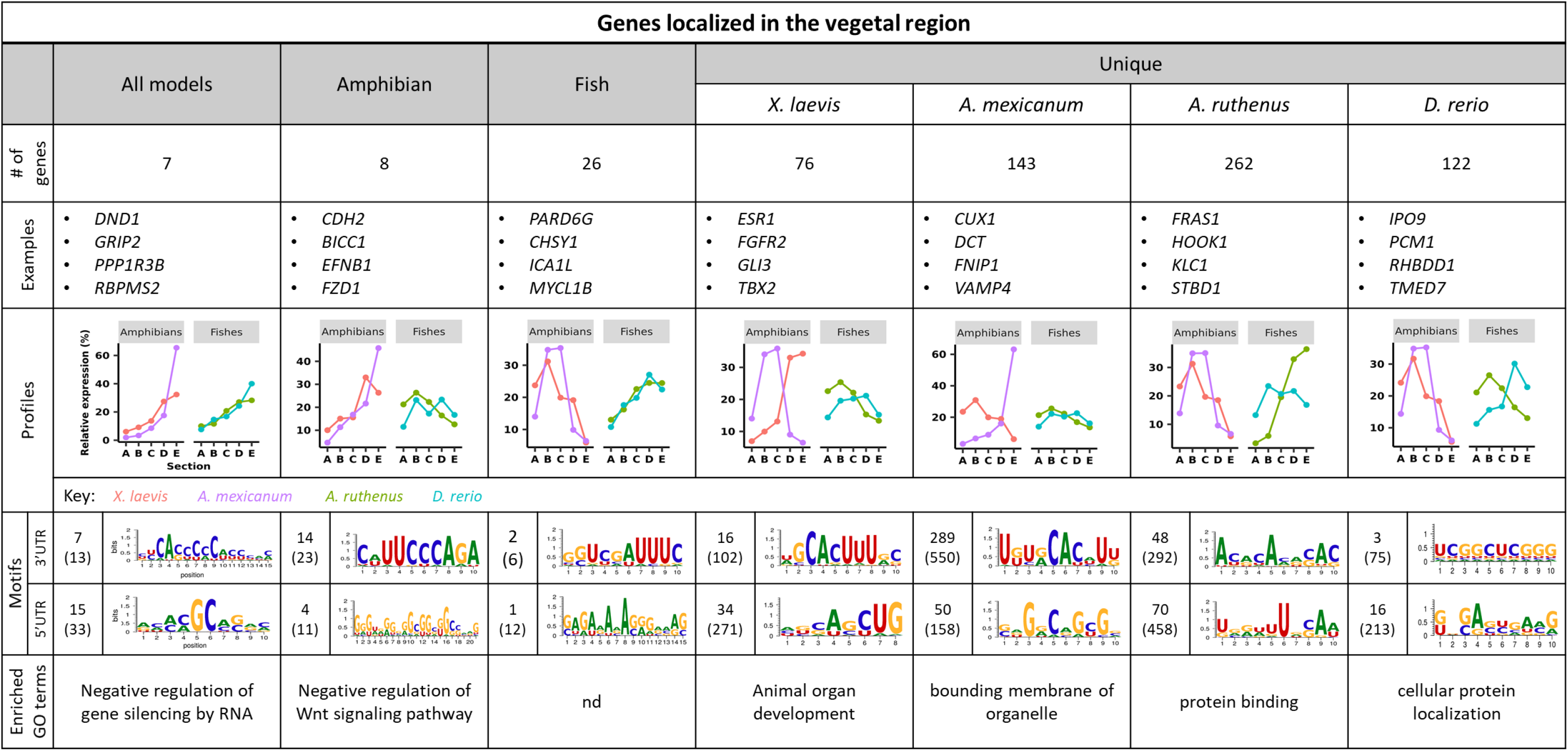
Number of extreme vegetal or vegetal localized genes conserved amongst the different models, as derived from dataset3. The motif numbers in brackets represent the motifs that were at least 2x more abundant in the given genes relative to the other sections, while the other motif number represents those that were also statistically significantly enriched. The motif image represents an example of one of the statistically significantly enriched 2x motifs as determined by AME. A representative of the enriched Gene Ontology (GO) term from the selected genes is shown in the last panel.

### Novel group of genes localized in the central region of eggs was revealed

In addition to the previously published groups of vegetal and animal genes, we identified a new group of genes that show central localization. The *A. mexicanum* and *A. ruthenus* had the most central DEGs, with 1454 and 138 respectively, while *D. rerio* and *X. laevis* had less at 24 and 6 respectively (dataset3, Fig. 2b; Supplemental file 2: Table S1, S5). There was little to no conservation of central genes amongst the models and no identical shared motifs within the UTRs (Fig. 2c, Fig. 4). Only *A. mexicanum* central genes displayed enriched GO terms related to organelle organization, the rest of the models did not have terms related to spatial arrangement or organization (Fig. 4). The complete list of enriched GO terms for the given categories is available in Supplemental file 2: Table S6. Analysis of the enriched GO terms, metabolic pathways, transcription factors and protein complexes for all the central DEGs, showed no overlap.

**Figure 4.**
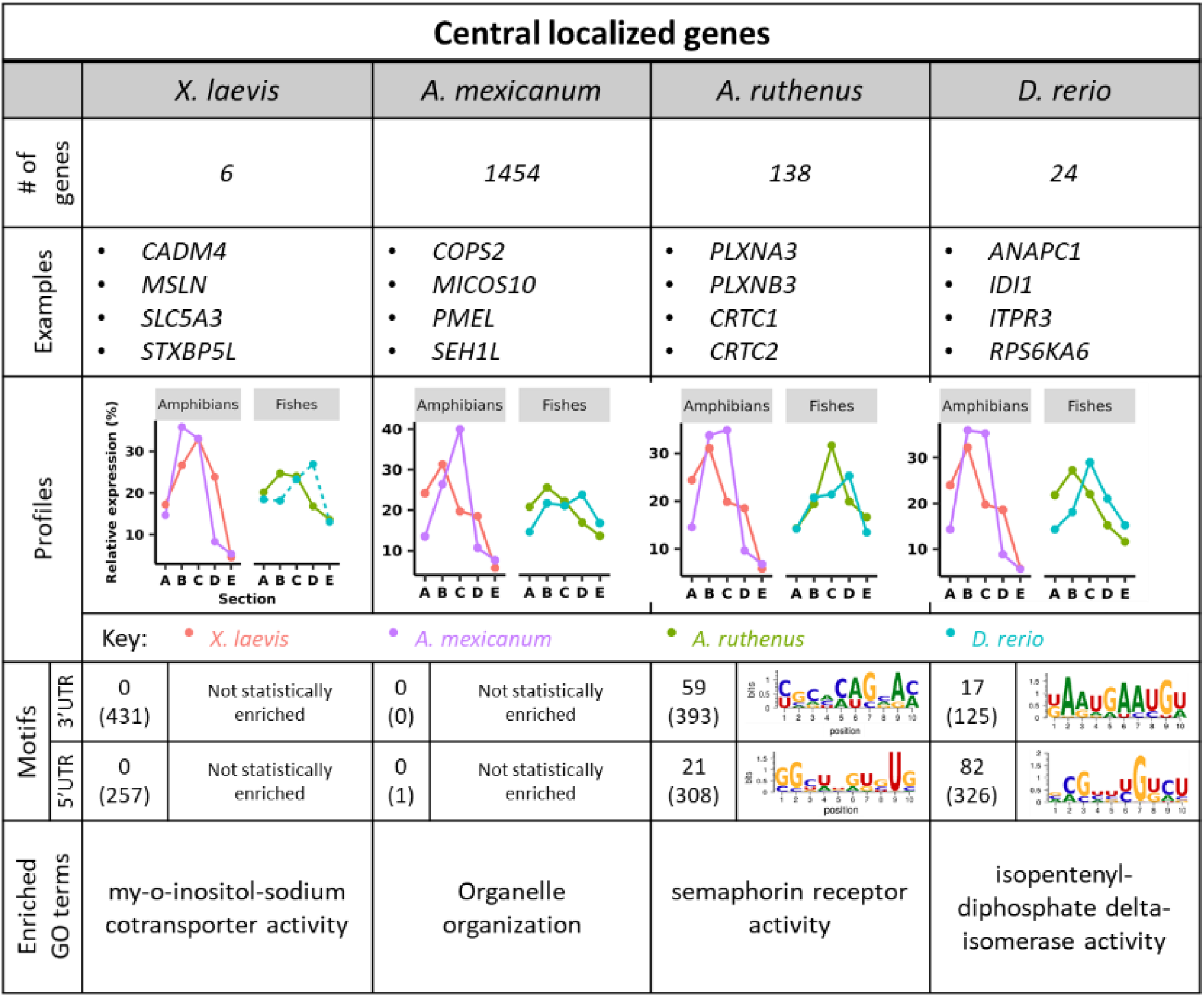
Number of central genes conserved amongst the different models, as derived from dataset3. The motif numbers in brackets represent the motifs that were at least 2x more abundant in the given genes relative to the other sections, while the other motif number represents those that were also statistically significantly enriched. The motif image represents an example of one of the statistically significantly enriched 2x motifs as determined by AME. A representative of the enriched Gene Ontology (GO) term from the selected genes is shown in the last panel.

### Animal localization showed higher level of conservation than vegetal and novel putative localization motifs

We found that the majority of DEGs were localized preferentially within the animal hemisphere and formed two distinct profiles, the extreme animal and animal profiles (Fig. 2b). We have already speculated within our previous work that most of animal mRNAs are still inside or around the nucleus region (Sindelka et al. 2018). However, in contrast the extreme animal mRNAs create a gradient with a maximum within the animal pole (segment A). The analyzed dataset3 contained uniquely 290 *X. laevis*, 206 *A. mexicanum*, 454 *A. ruthenus* and 99 *D. rerio* extreme animal or animal localized DEGs. Approximately 80 of these DEGs showed conservation of extreme animal/animal profiles among all species (Fig. 2c, Supplemental file 2: Table S1, S7). Extreme animal/animal DEGs found within all models showed enriched GO terms involved in cell cycle regulation and nuclear functions (Fig. 5). There were no enriched GO terms (probably due to the low number of genes with human symbols) for the DEGs unique to the amphibians, fish or each individual model, except for *A. ruthenus* which showed terms involving taste receptor binding (Fig. 5). The complete list of enriched GO terms for the given categories is available in Supplemental file 2: Table S8. Analysis of the enriched GO terms, metabolic pathways, regulatory motifs and protein complexes for all extreme animal and animal DEGs, showed a large overlap (3018) of similar terms between the models. The most interesting of these, appeared to be genes involved in the cell cycle regulation, RNA transport, cytoskeleton organization, the heart, cerebellum and skeletal system development and function.

**Figure 5.**
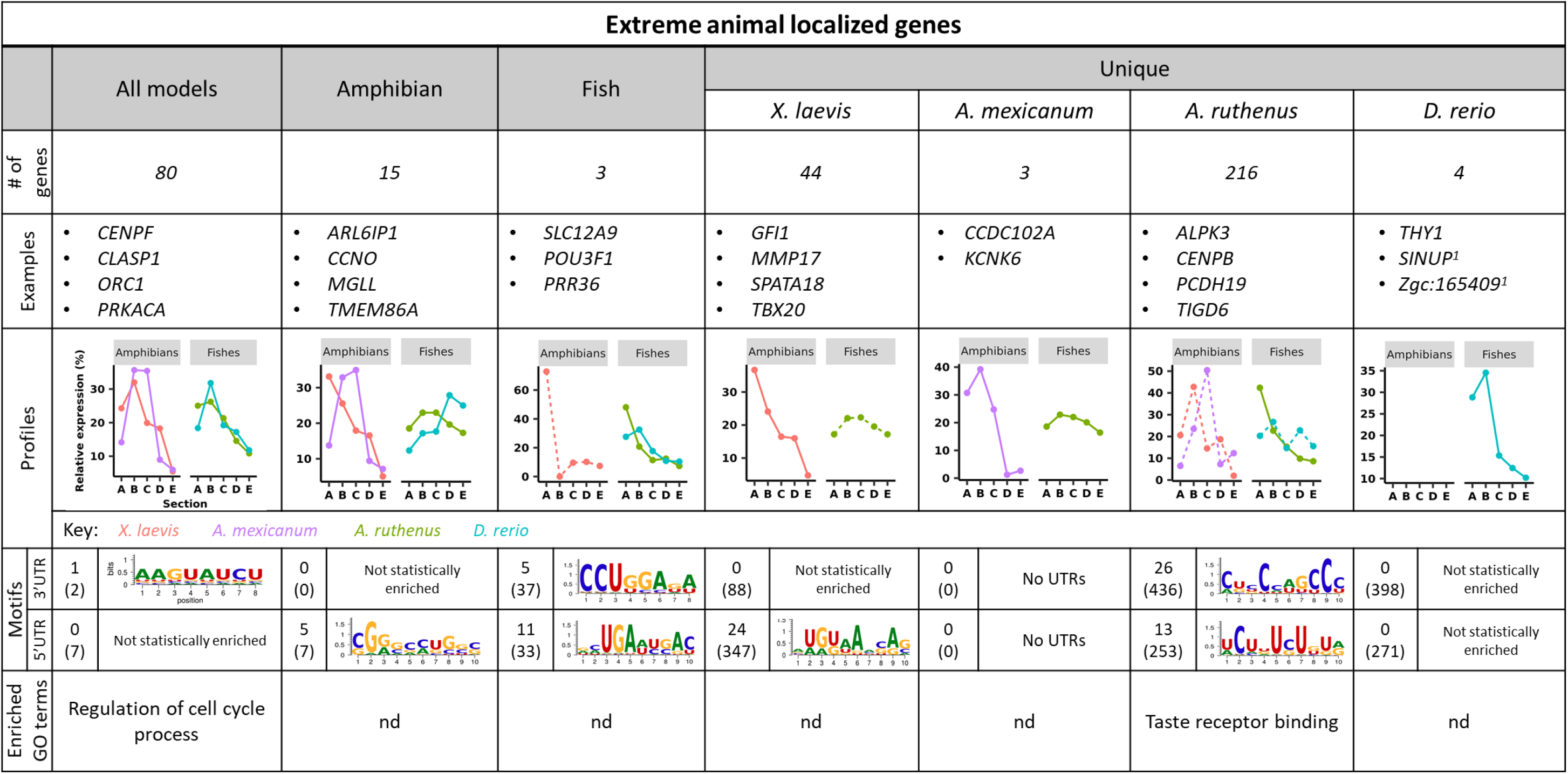
Number of extreme animal genes conserved amongst the different models, as derived from dataset3. The motif numbers in brackets represent the motifs that were at least 2x more abundant in the given genes relative to the other sections, while the other motif number represents those that were also statistically significantly enriched. The motif image represents an example of one of the statistically significantly enriched 2x motifs as determined by AME. A representative of the enriched Gene Ontology (GO) term from the selected genes is shown in the last panel. Orthologous animal DEGs to the extreme animal DEGs were included into the dataset. ^1^*Danio rerio* gene nomenclature.

### Motif analyses revealed good conservation of vegetal localization motifs and species dependent regulatory sequences

Motif analysis was performed using several independent tools to identify any putative localization sequences and any known regulatory sequences that were conserved within the 3’ and 5’UTR regions (datasets 2 and 3). Localization motifs with a CAC or GUU rich core were identified in the shared vegetal DEGs within all models (Fig. 3). The motifs unique to the amphibian specific vegetal DEGs were rich in Guanine or Cytosine, while another had a nucleotide core of UUCCCAG. The most significant fish specific motifs on the other hand, had core sequences with AUUUC or UUGAMGUG. There were many identified motifs, ranging from 75 to 550, that were identified within the vegetal DEGs specific for each model and most of them showed some variations around the core CAC. 5’UTR analysis of vegetal DEGs revealed similar motif numbers, with a CGC core being identified as shared amongst the four models. The amphibian motifs were Guanine/Cytosine rich or contained a GCG core, while the fish motifs were rich in Uracil residues. Despite the large number of central DEGs in the *A. mexicanum*, no statistically enriched motifs were detected in its UTRs, same as in *X. laevis* (Fig. 4). However, several putative motifs were identified within the fish. Analysis of extremely animal DEGs showed enriched motifs, however the majority of them did not pass our 2x enrichment and statistical significance requirements (Fig. 5). In general, we identified much fewer motifs compared to the vegetal group. In the 3’ UTR region, only two conserved motifs were found enriched in all models. The only statistically significant one: AAGUAUCU was present in less than 3.5% of the shared genes. Motifs with a conserved CCUGGA core were found in fishes and several enriched motifs were identified that were specific for each model, but only a few showed significance in *A. ruthenus*. In the 5’ UTR region, no statistically significant motifs were found for the animal genes shared amongst all models. The full list of the putative localization motifs that were statistically significantly more abundant and at least 2x more abundant relative to the other category can be found in Supplemental file 2: Tables S9-S11, while a heatmap of their distribution as derived from FIMO in Supplemental file 1: Fig. S4-S6.

Scan For Motifs was performed to identify any known regulatory sequences that might be responsible for certain behaviors of the RNAs (translation, stability, degradation etc.). We found an enrichment of several regulatory sequences within the 3’ UTR regions of the vegetal genes from the amphibians only. These sequences, Musashi binding element, GU-rich destabilization element UTRSite, CAG Element and Pseudoknot like structure are known to be involved in mRNA mediated decay, translational repression, transcriptional promoter and other diverse roles induced by secondary conformations. *A. mexicanum* contained about 18 more enriched regulatory motifs in their vegetal genes and they were also involved in the stability of the mRNA and polyadenylation. Only the *A. mexicanum* contained enriched protein binding sites (11) as determined from the RNA-Binding Protein DataBase. These binding proteins are usually associated with the degradation of the transcript, splicing, stimulation of transcription, mRNA export, bioaccumulation, or stabilization. There was no overlap in potential miRNA interactions among the four models. However, 16 miRNAs were associated with both of the amphibians’ vegetal genes, while a large quantity (1455) was associated with only the *A. mexicanum*. Analysis of the central group showed enrichment of two 3’ UTR regulatory sequences shared only within the amphibians: Musashi binding element and the secondary structure confirmation (pseudoknot like structure). Only *A. mexicanum* contained associated protein binding sites (12) while the amphibians were the only models to share sequences associated with miRNAs (18) and not the fishes. Analysis of the animal group found no shared enrichment of 3’ UTR regulatory sequences. Only two sequences (ARE motif, polyadenylation element) were found and each were associated with one of the fishes. There were no associated protein binding complexes that were significantly over-enriched for any of the animal genes. Only *D. rerio* (7) and *X. laevis* (2) had associated miRNA.

### Detailed analysis of known localized genes revealed surprising biological implications

We selected and analyzed in more details, genes whose functions during early embryogenesis and localization within the egg have already been well defined. We utilized RT-qPCR on independently prepared samples to either confirm our TOMO-Seq results or to determine the expression profiles of missing/highly variable known genes (usually low expressed transcription factors) (Fig. 6; Supplemental file 1: Fig. S7-S9). We also summarized the results of some of these known genes in *X. laevis* and *D. rerio* from other researchers for the validation of our RT-qPCR results (Supplemental file 2: Table S12). Below are four important categories such as germ plasm determinants, germ layer determinants, Wnt pathway responsible for dorsal/ventral specification and other interesting maternal genes (Wessely and De Robertis 2000; Claussen and Pieler 2004; Zearfoss et al. 2004; Yan et al. 2018).

**Figure 6.**
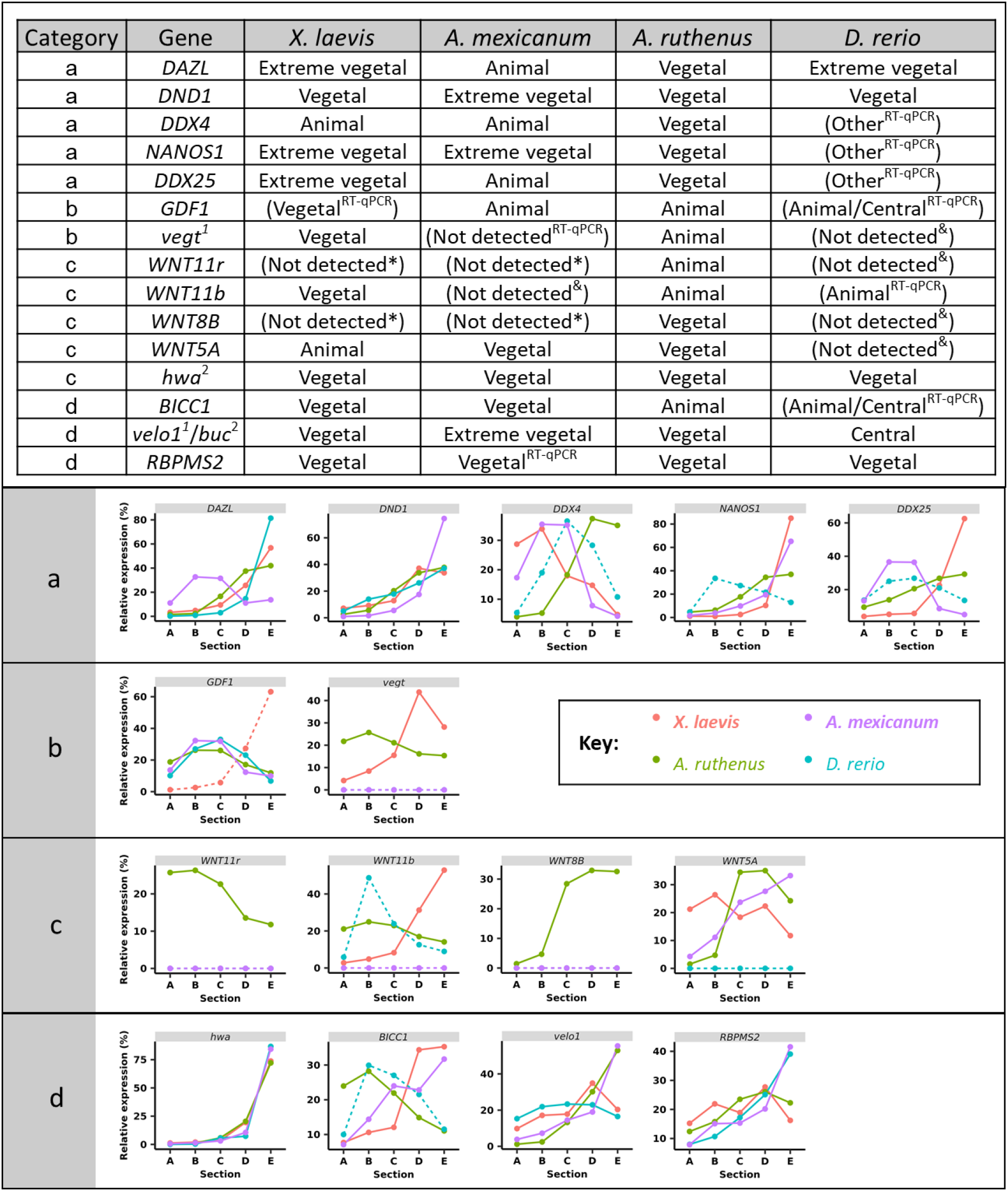
RNA localization of some essential genes within the egg of several species. Localization profiles for some key member genes belonging to (A) PGC markers, (B) Endodermal and mesodermal determinants, (C) wnt ligands, other (D) known *Xenopus laevis* vegetal genes. Genes in brackets and graphs with dashed lines indicate that the RNA-Seq data for the gene was not differentially expressed, failed the statistical analysis using DESeq or did not meet threshold criteria. Genes listed with “Other^RT-qPCR^” indicate that the RT-qPCR profiles were variable between the replicates. *Danio rerio GDF3* gene is an ortholog of *GDF1* gene. *velo^1^* (amphibians and *A. ruthenus*) and *buc^2^* (*D. rerio*) are orthologous genes. ^1^gene nomenclature for *X. laevis;* ^2^gene nomenclature for *D. rerio*; *primers worked on later stages; ^&^not detected even at later stages.

The germ plasm determinant category was selected based on literature. These comprised of genes such as *DND1*, *DDX4* (also called *vasa*/*xvlg1*), *DAZL*, *DDX25* (previously *deadsouth*) and *NANOS1* which contribute to the germ plasm and PGCs migration and survival (Weidinger et al. 2003; Theusch et al. 2006; Kosaka et al. 2007; Flachsova et al. 2013; Lasko 2013). We found a perfect conservation of vegetal localization for *DND1* and *GRIP2* PGCs markers across all models. *DAZL* mRNAs were localized vegetally for *X. laevis, D. rerio* and *A. ruthenus*, but animally localized in *A. mexicanum* egg. *DDX4* mRNAs were detected vegetally in *A. ruthenus*, but animally or showing another location in the remaining models. mRNAs coding *NANOS1* were found in the vegetal region in amphibians and *A. ruthenus* but at variable positions (ranging from animal to vegetal) in *D. rerio. DDX25* mRNAs were localized in the vegetal region only for the *X. laevis* and *A. ruthenus* eggs.

The majority of published information about endoderm and mesoderm specification comes from research done using the *X. laevis* model and has led to the characterization of two well-known transcription factors *GDF1* (in *D. rerio* homolog *GDF3*) and *VEGT*. Our TOMO-Seq and RT-qPCR revealed vegetal localization only in *X. laevis* and animal localization or no detection in the remaining species. The *VEGT D. rerio* homolog was not detectable even by RT-qPCR in either the eggs or at the blastula stage.

The Wnt pathway is responsible for the dorsal/ventral axis specification. Currently, most of our knowledge about this pathway has been based on studies done on the *X. laevis* and *D. rerio* models. These studies have shown that the key ligands are *WNT11* (*WNT11r* and *WNT11b* paralogs), *WNT8B* for the canonical and *WNT5A* for the noncanonical pathways. Wnt ligands are usually very lowly expressed and therefore their quantification is challenging. We identified vegetally localized *WNT11b* in *X. laevis*, while vegetally localized *WNT8B* and *WNT5A* in *A. ruthenus*. The remaining genes such as *AXINS*, *GSK3B* and *CTNNB1* were not detected or were animally localized. A recent study (Yan *et al.,* 2018) identified another member of the Wnt pathway called *huluwa* (*hwa*) in *D. rerio*. We manually blasted and annotated its potential homologous genes in the remaining species and found that it is vegetally localized in all models and can be added to the vegetally conserved group.

We also validated the localization profiles for other developmentally important maternal genes such as *RBPMS2*, *BICC1* and *VELO1/BUC* and found vegetal conservation for only *RBPMS2* (previously *hermes*). *BICC1* is vegetally localized in amphibians and animal in fish models. *VELO1* is localized vegetally in *X. laevis*, *A. mexicanum* and *A. ruthenus*, while its homolog *BUC* is instead localized centrally in *D. rerio*.

## Discussion

Localization of mRNA molecules inside the tissue and even inside the single cell such as the oocyte and zygote, leads to asymmetrical translation. This is a well-known phenomenon in the fields of cellular and developmental biology (Marlow 2010). *Xenopus laevis* and *D. rerio* belong to the most popular vertebrate animal models used for studying development and many important genes involved in this essential process were discovered by using these models during the last few decades (Kloc and Etkin 1995; Yoon et al. 1997). Our goals here have been to elucidate RNA localization at a global scale and to compare the localization profiles of homologous genes with a focus on the evolutionary perspective. Four animal models covering two main branches of evolution, *Amphibia* and *Actinopterygii* were analyzed using the same experimental workflow. Even though some of them belong to the same class, there are several differences in their early development (type of fertilization, cleavage pattern, germ layer formation, PGC specification, etc. - reviewed in Introduction).

### Overcoming technical challenges

In our work we observed between 5-90% of DEGs within the four models, relative to their assessed transcriptomes. We annotated between 49-89% of the DEGs using known and *de novo* prepared transcriptomes. However, the quality of data is dependent on both the quality of the sequences and its annotation. Here, we had to overcome the first obstacle because *A. ruthenus* and *A. mexicanum* genome annotation is far from perfect. However, by validating the homology manually and utilizing strict criteria for the data quality control, we were able to identify the majority of the maternal pool within these models. Another obstacle was the identification of DEGs, determination of localization profiles and calculation of reproducibility among biological replicates. In our previous work we identified 4 main groups of localization profiles in *X. laevis* based on very simple parameters. We were able to categorize >99% of the identified maternal DEGs. Surprisingly, the same parameters were applicable on the other three datasets, and we were able to cluster > 80% of DEGs into groups. We added a new group, called central, which is primarily absent in *X. laevis*.

Of course, the number of identified genes was directly proportional to the quantity of available material and quality of TOMO-Seq data. *Danio rerio* eggs are smaller and contain at least 5-fold lower concentration of total RNA compared to the other models. Unfortunately, because the eggs are transparent and become hidden in the medium after freezing, external markers such as a surrounding blue dye or stained bead must be added to the cryomedia. Cryosectioning must be performed by using just a single egg, with its correct orientation validated using RT-qPCR of known vegetal and animal genes. For this reason, the starting material for library preparation was limited and we found low read numbers for many maternal genes and a high level of variability among replicates. Therefore, the total number of genes, which passed our quality criteria is much lower than for the other models. After filtering, TOMO-Seq results from *X. laevis*, *A. mexicanum* and *A. ruthenus* provided reproducible quantification of > 80% of known transcriptomes and we can speculate that this is probably covering nearly complete maternal mRNA pools. However, we observed an absence or very low counts (resulting in non-DEG evaluation) for several important transcription factors such as *GDF1* and *WNTs*, and therefore additional RT-qPCR was performed to improve the quality and interpretation of our results.

### Evolutionary conservation of localization profiles is modest, despite good motif conservation

Our analyses started with the hypothesis that the complete extra uterine development of our models without any additional environment stimuli from the mother, must be propagated in a predetermined organization of biological molecules formed during oogenesis. Our results support this theory and revealed that the great majority of identified mRNAs are asymmetrically localized and create just 5 distinct localization profiles (extreme animal, animal, central, vegetal and extreme vegetal). This is contradictory to the traditional view, where most of mRNAs are expected to be ubiquitous and evenly distributed with the exception of just a few asymmetrically localized mRNAs coding genes responsible for developmental plan specification. In general, we observed all localization groups in all our chosen animal models, however the proportions were different. In most of the models, the majority of the localized mRNAs form the **animal** profile with enrichment in the second section, which corresponds to the region with germinal vesicle and later nucleus position. Gene function analysis suggest that these mRNAs are required for activities connected with nuclear function, cell division and other housekeeping processes. In our previous work we speculated that these mRNAs do not undergo active transportation and that the animal profile is created by diffusion (Sindelka et al. 2018). We identified an additional sub-profile within the animal hemisphere that was characterized by a steeper gradient from the animal to vegetal pole, and we called this profile **extreme animal**. We found the extreme animal profile in all models, but there were differences in the gene numbers and profile shapes. *Acipenser ruthenus* and *X. laevis* had a much higher proportion of extreme animal genes compared to the remaining models and *A. ruthenus* showed much steeper animal gradients compared to *X. laevis*. Comparison of animally enriched genes with focus mainly on extreme animal genes revealed a large number of conserved genes amongst our models. We also identified several enriched motifs in the 3’ and 5’UTRs, which could be considered as putative localization motifs for animal localization. Interestingly, there were differences in the shared and species specific motifs and the GO terms. The third group of genes was called **central**. *Xenopus laevis* showed a very low proportion of central genes compared to the other groups, while in contrast *A. mexicanum* showed quite a large proportion of centrally localized mRNAs. Overlap of central genes between models was poor and we did not find any conserved gene or pathway. In addition, GO term analysis showed poor results. The last two groups are called **vegetal** and **extreme vegetal** based on the steepness of their gradients. Many *X. laevis* and *D. rerio* vegetal genes have already been extensively studied and their role in development is well known. Surprisingly, comparative analysis showed poor conservation and revealed only 7 conserved genes. Most of these genes have already been studied and are known to be involved in PGCs migration and survival (*DND1*, *GRIP2*) and also serving as RNA-binding proteins (*RBPMS2*) (Weidinger et al. 2003; Kirilenko et al. 2008). An interesting group of 15 genes showed vegetal localization in three models only, while in the fourth model they were animally/variably localized. Among these genes belong *DAZL*, *VELO1/BUC, SLAIN1* and *SULF1*. Interestingly there are several genes that are selectively vegetally localized in either amphibians or fishes. As an example, we can find vegetal *BICC1* in amphibians, but with other profiles or even absent in fishes. In contrast, *CELF1* and *TGFA* genes are vegetal in fishes but different in amphibians. An interesting category are *SLC* (solute carrier family) genes, where some members show amphibian/fish specific vegetal localization. There are also species specific vegetal genes such as *VEGT* and *Xpat (also called pgat*) in *X. laevis*.

We selected several known important maternal genes for thorough analysis. One of the key maternal factors are the primordial germ cell determinants. We identified many genes such as *DND1*, *DAZL*, *NANOS1* to be preferentially vegetally localized (in exception of *DAZL* in *A. mexicanum* and *NANOS1* in *D. rerio*), however others such as *DDX4* and *DDX25* showed little localization conservation. This contrasting localization of *DAZL* in *A. mexicanum* has already been observed using other approaches (Bachvarova et al. 2004) and coincides with the differently proposed mechanism of its PGC formation. This suggest that several mechanisms of PGC formation exist and that it is also species dependent. *Xenopus laevis* and *A. ruthenus* are considered as the most similar species of this study in respect to PGC development. They possess PGC specification by preformation of PGC precursors at the vegetal pole and they both demonstrate holoblastic cleavage patterns during early embryonic development. *Danio rerio* specifies their PGC also by preformation, however its cleavage pattern is meroblastic, thus their PGCs are formed at the marginal region of the blastodisc. *Ambystoma mexicanum* has a completely different PGC specification mechanism using epigenesis. The PGCs are induced from pluripotent cells by signals from the surrounding somatic tissues. *VEGT* and *GDF1* genes are responsible for ectoderm and mesoderm specification, however we found very little conservation in their profiles, which also suggest an independent mechanisms of germ layer formation or presence of other substituting factors. A similar interpretation can be made for Wnt signaling. Vegetal localization is species dependent (*WNT11b* is vegetal only in *X. laevis*) and usually other members of the Wnt pathway are localized in the animal profiles. Interestingly, a gene interacting with the Wnt pathway during dorsal specification called *hwa*, is vegetally localized in all models.

In contrast to the low conservation at the gene level for vegetal localization, we revealed a good conservation of putative localization motifs. CAC-rich motif was conserved among our models, and we found many species specific CAC-rich variants. In addition, we identified several other conserved and species specific putative motifs. An interesting observation was the enrichment of known regulatory sequences in the amphibian’s vegetal genes, while the absence of this enrichment in fishes indicated a potentially later activity for their vegetal genes. We can speculate that the determination of the fish germ layers requires vegetal - central RNA transportation after fertilization and that the fish vegetal RNAs are used at the later developmental stages (e.g yolk stream in *D. rerio*) (Theusch et al. 2006; Kosaka et al. 2007; Houston 2013). In contrast to the vegetal motifs, the conservation of the extremely animal and central motifs was lower, and we found mainly species specific variants. Also, enrichment of the regulatory sequences was limited and species dependent and suggest a lower translational activity of animal RNAs.

In summary, our results indicate that processes such as germ layer formation, primordial germ cell specification and determination of body axes are much more complex than the expectations outlined in the current literatures. Even though we believe that mRNA localization is a driving force of animal-vegetal asymmetry, specification of other body axes is probably based also on other factors or on a combination of various molecules. Poor evolutionary conservation of the localization of mRNAs coding important transcription factors, especially in the vegetal region, suggest that the mechanisms are species dependent. In contrast to poor gene localization, there is a good correlation in putative localization motifs present, with the CAC-rich motif probably being the key motif. However, there are also species specific CAC motif modifications. We can speculate that these motif variants are important for the species specific RNA-binding role of proteins responsible for RNA accumulation, preservation and transportation.

## Material and Methods

### Ethics approval

All experimental procedures involving model organisms were carried out in accordance with the Czech Law 246/1992 on animal welfare. *Acipenser ruthenus* and *D. rerio* animals are from colonies of the Research Institute of Fish Culture and Hydrobiology in Vodnany, Czech Republic and protocols were reviewed by the Animal Research Committee of the Faculty of Fisheries and Protection of Waters, South Bohemian Research Center of Aquaculture and Biodiversity of Hydrocenoses, Research Institute of Fish Culture and Hydrobiology, Vodnany, Czech Republic. The *X. laevis* animals were from the colony of the Institute of Biotechnology and protocols were approved by the animal committee of the Czech Academy of Sciences. *Ambystoma mexicanum* animals were from the colony of the Department of Zoology, Faculty of Science, Charles University, Prague, Czech Republic and protocols were approved by the Faculty of Sciences of the Charles University.

### Model organisms

*Xenopus laevis* females were stimulated with 500 U of human gonadotropin, left overnight at 18°C, and the eggs produced the following day were collected after jelly coat removal. Twenty eggs were prepared together in the sectioning block. *Ambystoma mexicanum* male and female were incubated overnight together in an aquarium at 15°C to naturally stimulate egg maturation and fertilization. Laid eggs/embryos were collected and visually inspected. *Acipenser ruthenus* ovulation in the females was stimulated by intramuscular injection of carp pituitary extract in two doses. The first was given at 36 h before egg stripping (0.5 mg/kg of body weight) and the second at 24 h before egg stripping (4.5 mg/kg of body weight). Ovulated eggs were sampled using the microsurgical incision of the oviducts in water as described by Podushka (1999). *Danio rerio* eggs were obtained from females and washed for 3 minutes in water to display the blastodisc, which served as a selection criterion of functional eggs and guide for their orientation. Trypan blue staining solution at a concentration of 0.001 % was added to the eggs, after which the individual eggs were transferred to the sectioning medium and oriented before freezing.

Samples were prepared in at least biological duplicates using two independent experiments and different females (in total >4 samples). Eggs (*X. laevis*, *A. ruthenus* and *D. rerio*) or potentially one-cell stage embryos (*A. mexicanum*) were embedded in Tissue-Tek O.C.T. Compound, oriented along the animal-vegetal axis (animal pole positioned at the top) using delicate forceps and immediately frozen on dry ice and stored at −80°C.

### Sample preparation

Samples were incubated for 10 minutes in the cryostat chamber (−20°C) and then cut into 30 *μ*m slices along the animal–vegetal axis. Slices were consequentially pooled into five tubes with the same number of slices per tube. Tubes were then labelled to correspond to the relevant segments of the oocyte: (A) animal cap - (E) vegetal cap. Total RNA (*X. laevis* and *A. mexicanum*) was extracted using TriReagent extraction and LiCl precipitation (Sigma) (details in (Sindelka et al. 2018)). The *A. ruthenus* samples were extracted using Qiagen (Minikit + LiCl precipitation) according to the manufacturer’s instructions. Given the smaller size of *D. rerio* eggs, Qiagen (Microkit), column-based isolation was used for its RNA extraction. The concentration of RNA was measured using a spectrophotometer (Nanodrop 2000, Thermo Scientific), and the quality of RNA was assessed using a Fragment Analyzer (AATI, Standard Sensitivity RNA analysis kit, DNF-471). No signs of RNA degradation were observed. Absence of inhibitors and the precision of the orientation of the embedded egg were tested using RT-qPCR quantification of the RNA spike (TATAA Biocenter) and known localized marker genes respectively.

The cDNA was prepared using total RNA (*X. laevis -* 20 ng, *A. mexicanum* – 30 ng, *A. ruthenus –* 40 ng and *D. rerio* – 20ng), 0.5 *μ*l of oligo dT and random hexamers (50 *μ*M each), 0.5 *μ*l of dNTPs (10mM each) and 0.5 *μ*l of RNA spike (TATAA Universal RNA Spike, TATAA Biocenter), which were mixed with RNAse free water to a final volume 6.5 *μ*l. Samples were incubated for 5 minutes at 75°C, followed by 20 seconds at 25°C and cooling to 4°C. In the second step, 0.5 *μ*l of SuperScript III Reverse Transcriptase (Invitrogen), 0.5 *μ*l of recombinant ribonuclease inhibitor (RNaseOUT, Invitrogen), 0.5 *μ*l of 0.1 M DTT (Invitrogen), and 2 *μ*l of 5 × First strand synthesis buffer (Invitrogen) were added and incubated: 5 minutes at 25°C, 60 minutes at 50°C, 15 minutes at 55°C and 15 minutes at 75°C. Obtained cDNAs were diluted to a final volume of 100 *μ*l and stored at −20°C.

### Primer design and quantitative PCR

Primer assays of selected maternal genes were designed using NCBI Primer-Blast (https://www.ncbi.nlm.nih.gov/tools/primer-blast/) (NCBI). Expected amplicon length was set to 70-232 bp and Tm to 60 °C. Primer sequences are available in Supplemental file 2: Table S13. The RT-qPCR reaction contained 3.5 *μ*l of TATAA SYBR Grand Master Mix, 0.29 *μ*l of forward and reverse primers mix (mixture 1:1, 10 *μ*l each), 2 *μ*l of cDNA and 1.21 *μ*l of RNase-free water in 7 *μ*l final volume. RT-qPCR was performed using the CFX384 Real-Time system (BioRad) with conditions: initial denaturation at 95°C for 3 minutes, 45 repeats of denaturation at 95°C for 15 seconds, annealing at 60°C for 20 seconds and elongation at 72°C for 20 seconds. Melting curve analysis was performed after to test reaction specificity and only one product was detected for all assays. Only samples with continuous gradient profiles of the marker genes were selected for library preparation.

### Library preparation

Given that the collection of samples from all four animal models was done at different times, we used a variety of depletion and library preparation kits. Quality and performance of each kit was tested beforehand, and we selected the best option. Details about the RNA quantity, depletion, library preparation kits and RNA sequencing are presented in Supplemental file 2: Table S14. Library qualities were assessed using the Fragment Analyzer (AATI, NGS High Sensitivity kit (DNF-474) and the concentration was determined by the Qubit 4 Fluorometer (ThermoFisher Scientific). Equimolar library pools were prepared and sequenced.

### RNA-Seq workflow

A brief summary schematic of the RNASeq data analysis can be found in Supplemental file 1: Fig. S10. Adaptor sequences and low-quality reads were filtered out using TrimmomaticPE (v. 0.36) (Bolger et al. 2014) using the parameters as outlined in Supplemental file 2: Table S15 and “LEADING:3 TRAILING:3 SLIDINGWINDOW:4:15 MINLEN:36”. SortMeRNA (v. 2.1b) (Kopylova et al. 2012) was used to remove mtRNA reads (Supplemental file 2: Table S15) and any remaining rRNA reads. Next, reads were aligned against the model’s respective genome (Supplemental file 2: Table S15) (*X. laevis* (Xenbase), *D. rerio* (Ensembl)) using STAR (Dobin et al. 2013) and counted using htseq-count or directly counted against the model’s transcriptome (*A. ruthenus* (in-house), *A. mexicanum* (Axolotl-omics)) using kallisto (v. 0.43.1) (Bray et al. 2016). The data were deposited in the National Center for Biotechnology Information’s Gene Expression Omnibus (GEO) (Supplemental file 2: Table S14).

### Data processing (normalization, clustering, annotation)

The gene counts across each section for each model were normalized using the NormQ method (Naraine et al. 2020), followed by differential gene expression analysis using DESeq2 (v. 1.24.0) (Love et al. 2014). The list of marker genes used for the normalization can be found in Supplemental file 2: Table S16. Due to the presence of several lowly expressed variants in the *A. mexicanum*, only genes with a raw (before normalization) gene count greater than 30 copies in any sample were analysed. The design of the experiment followed a “∼replicate + position” setup, while differential analysis was carried out using DESeq2’s default parameters, with the use of the Likelihood-Ratio-Test using a reduced model of “∼replicate” and a padj value cut-off of 0.1. The localization profile of each gene consisted of five categories: extreme animal, animal, central, vegetal and extreme vegetal. The clustering was based on the criteria as defined from our previous study, with the inclusion of the additional central profile, and can be found in Supplemental file 2: Table S17 (Sindelka et al. 2018). Genes whose profiles did not fit into these defined five localization categories were classified as “undefined”.

We created three different datasets applicable for each downstream analysis. Dataset1 (Supplemental file 2: Table S1) comprised of all the differentially expressed genes (DEGs – genes with reliable expression and uneven distribution of RNAs) from each model and was used for a less stringent ortholog and localization comparative analysis, paralog analysis (by matching gene symbols) within the same model and comparative GO analysis. Dataset2 (Supplemental file 2: Table S1) comprised of reproducible DEGs with well-defined profiles and was used for motif enrichment analysis and detection of known motifs. The parameters for inclusion into the dataset2 are outlined in Supplemental file 2: Table S18. Dataset3 (Supplemental file 2: Table S1) comprised of a smaller subset of curated DEGs with well defined, reproducible profiles followed by additional annotation analysis, and was used to carry out a more comprehensive ortholog comparative analysis between the models.

Dataset1 was used to analyse for global conservation of the maternal transcripts, contrasting locations in paralogs and comparison of enriched GO terms. Comparisons using dataset1 were made between the models on the conservation of all annotatable DEGs, the annotatable animally localized (extreme animal/animal) DEGs, the annotatable centrally localized DEGs and the annotatable vegetally localized (extreme vegetal/vegetal) DEGs. Potential paralogous DEGs within the same models that showed very contrasting localization profiles (extreme animal/animal versus extreme vegetal/vegetal) were also identified by matching genes of the same symbols. The dataset2 comprised of two animal profiles, one central and two vegetal profiles and was used to analyse for known motifs, motif distribution and motif enrichment. Dataset3 was used to do a more in-depth verification of the conservation of the maternal DEGs. Genes that were extremely animal, central and vegetal (extreme vegetal/vegetal) were targeted. Originally, genes that were enriched within Section A were targeted for the analysis of the extremely animal genes. However, given the small number of extreme animal genes found in *A. mexicanum* and *D. rerio*, extreme animal and animal genes enriched in section A and section B were included into the extreme animal dataset to increase the power of the downstream analysis. The later annotation of these genes resulted in the detection of orthologs that were located in the animal categories. These gene were added to the dataset to produce the final dataset. The central genes analyzed were the same as those from the dataset2 while the vegetal genes originally comprised of all the extreme vegetal and vegetal genes from dataset1. These vegetal genes were then manually analyzed to select only reproducible and well-defined profiles. The dataset3 was used for *de novo* motif analysis and ortholog and localization comparisons. All overlapping genes from dataset3 that shared a localization profile with at least one other model organism were verified for correct ortholog matching and location placement by checking the replicate profiles and the nucleotide (e-value < 0.001) and/or protein alignment (blast v. 2.2.31) (Camacho et al. 2009) relative to the respective human reference or relative to a given model.

The organisms’ gene identifiers were mapped to a most probable *Homo sapiens* ortholog gene symbol by extracting its official *H. sapiens* gene ortholog, comparing its gene symbol to those from *H. sapiens* or comparing its protein sequence. The official ortholog mapping between *X. laevis* genes and *H. sapiens* genes were extracted from the Xenbase (http://www.xenbase.org/, RRID:SCR_003280) (04/10/18) (Xenbase; Fortriede et al. 2020). The official *H. sapiens* orthologs for *D. rerio* were extracted using the biomaRt (v. 2.40.5) R package using reference data from www.ensembl.org (∼02/08/19) and dataset from ENSEMBL_MART_ENSEMBL (Durinck et al. 2005, 2009). In the event of multiple *H. sapiens* gene ortholog symbols mapping to the *D. rerio* gene, the symbol that matched the query gene symbol was used, or in the absence of a match the first mapped symbol was used. The annotation of the *A. ruthenus de novo* transcriptome was done using Trinotate (v. 3.0.1) (Bryant et al. 2017) while the annotation for *A. mexicanum* was extracted from the data provided from Nowoshilow et al. 2018. In all four models, gene symbols that could not be officially obtained were mapped to its most likely *H. sapiens* gene symbol by matching the exact name of the query gene against all known *H. sapiens* gene symbols from the *H. sapiens* annotation R package org.Hs.eg.db (v. 3.8.2) (Carlson 2019). The ortholog matching was supplemented with the reciprocal best alignment heuristic tool Proteinortho (v. 6.0.9) along with DIAMOND (v. 0.9.35) (Lechner et al. 2011; Buchfink et al. 2014). This software was used to map the protein orthology between all four models along with the inclusion of the *H. sapiens* proteome (GRCh38.p13) as reference. Genes with missing *H. sapiens* symbols were appended with a *H. sapiens* symbol if they formed ortholog clusters with a given *H. sapiens* protein. In the absence of a known symbol, the ortholog cluster number was used as an identifier to trace the ortholog across the models. The data was also used to identify potential paralogs within the same model’s transcriptome. The gene from each organism was also mapped to all orthologs from the other models. Mapping was based first on the official *H. sapiens* ortholog symbol and gene symbol matches, and then to the ortholog cluster number assigned by Proteinortho.

### Motif analysis

Conserved motifs present within the Untranslated Regions (UTRs) of the significantly enriched genes that shared localization across all models, between amphibians, between fishes, or unique to each individual model (referred to as genes of interest) for the dataset3 (extreme animal, vegetal (vegetal/extreme vegetal) and central genes) were analysed for the presence of potential localization/zip-code motifs or known cis-regulatory motifs that may be responsible for their location or stability and translational efficiency. Several motif detection software were utilized and consisted of MEME (v. 4.11.2) (Bailey and Elkan 1994) using anr (motif width = 6 to 25, e-value < 0.05, max. number of motifs = 10), zoop (with and without a prior; e-value < 0.05, max. number of motifs = 10); DREME (e-value < 0.05) (v. 4.11.2) (Bailey 2011); homer2 denovo (v. 4.8.3) (Heinz et al. 2010) (motif widths = 6, 8, 10, 12, max. number of motifs = 100, fullMask); BoBro (v. 2.0) (Li et al. 2011), Weeder2 (v. 2.0) (motif widths = 6, 8, 10, threshold = 50, p scoring = 25) (Pavesi et al. 2004; Zambelli et al. 2014) and BioProspector (v. 2004 release) (Liu et al. 2001) (motif widths = 6, 8, 10, max. number of motifs = 10). All detected motifs were converted to the standard MEME motif format. Where possible the contrasting profiles or contrasting orthologous genes were used as negative controls during motif analysis. The MotifComparison (v. 3.2.2) (Claeys et al. 2012) tool (algorithm = KL; threshold = 0.001) was used to remove redundant (score = 0; shiftm-d = 0; shiftd-m = 0) motifs that were detected from the same location. Enrichment analysis of the motifs was done against the subset of the genes within dataset2.

FIMO (v. 4.11.2) (thresh 1e-4) (Grant et al. 2011) was used to scan the occurrence of each of the detected *de novo* motifs within the list of localized genes from dataset2. Motifs were then filtered to select for those that were at least 2x more abundance in the section of interest/genes of interest/model of interest versus the other localization categories/models. AME (v. 4.11.2) (--pvalue-report-threshold 0.05) (McLeay and Bailey 2010) was then used to determine whether these detected motifs were statistically enriched solely within the given localization category (dataset2) or set of genes of interest. TOMTOM (v. 4.11.2) (Gupta et al. 2007) was used to detect the most probable known relative of the detected motifs using the database Ray2013_rbp_All_Species and e-value cut-off of 0.01. Known interacting cis-regulatory motifs, protein binding complexes and miRNA were detected within the 3’UTRs of the subset of localized genes (dataset2) using Scan For Motif (access date: 24/10/20) (Biswas and Brown 2014). The detected known motifs were then analysed for significant enrichment within a given section using the one-tail Fisher Exact Test (p < 0.01).

### Gene ontology and Pathway analysis

Significant enrichment of Gene Ontologies, biological pathways, regulatory motifs and protein complexes were analysed using the R package, gprofiler2 (v. 0.2.0) (Raudvere et al. 2019), using the default parameters except for correction method = “g_SCS/fdr”, user threshold = “0.05”, domain scope = “annotated”, background organism = *H. sapiens*/*X. tropicalis*/*D. rerio* or custom background from GRCh38.p13. Genes of interest included those specific to all models, to amphibians, to fishes and to each model for the different localization categories that were analysed during the cross species ortholog analysis (dataset3). The total animal (extreme animal and animal), vegetal (extreme vegetal and vegetal) and central DEGs (dataset1) from each model were also analysed. The overlap of the enriched terms within the same categories of the different models was then analysed.

## Data Availability

All raw and processed sequencing data generated in this study have been submitted to the National Center for Biotechnology Information’s Gene Expression Omnibus (GEO) database, at https://www.ncbi.nlm.nih.gov/geo/, and can be accessed with the GEO deposition number: GSE104848 (*Xenopus laevis*), GSE166916 (*Ambystoma mexicanum*), GSE125819 (*Acipenser ruthenus*) and GSE166917 (*Danio rerio*).

## Acknowledgements

We thank our colleagues: O. Smolik, S. Tomankova, K. Pocherniaieva, M. Valihrachova and other members of participating laboratories for assistance with sample preparation.

## Author Contributions

VI, RS and RN wrote the manuscript and formulated methods. PA performed library preparation for RNASeq analysis and initial Bioinformatics analysis. RN performed Bioinformatics analysis. VI prepared cryosection of the eggs, RNA isolation and RT-qPCR. RF, VS, MP prepared eggs of different models for sectioning and were involved in result interpretation. All authors reviewed the manuscript.

## Additional Files

### Supplementary file 1

**Supplementary file 1: Figure S1.**
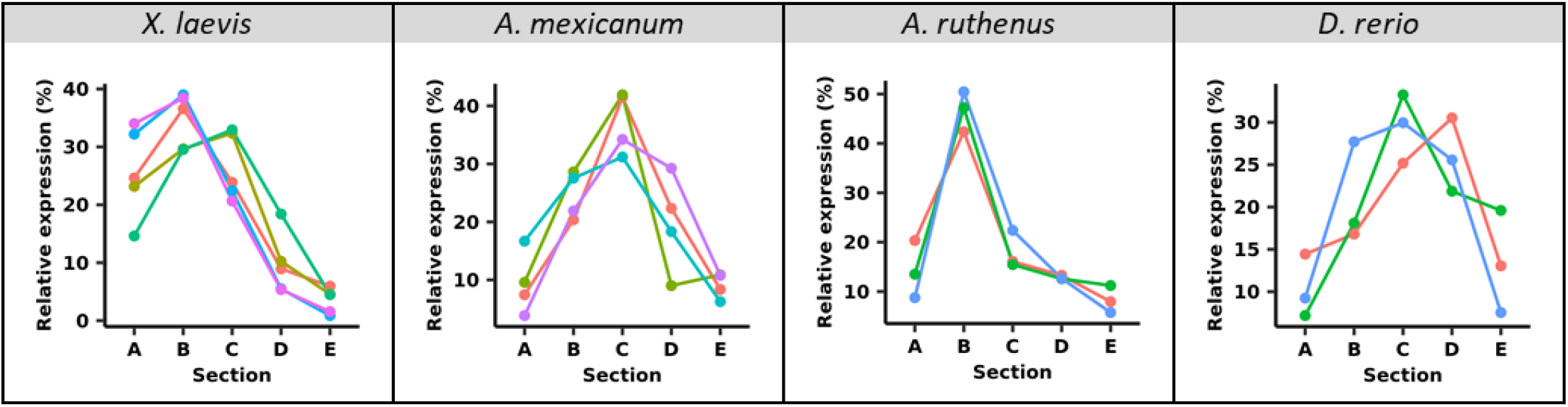
Relative RNA concentration extracted from each section of the egg. Each separate line represents a different replicate.

**Supplementary file 1: Figure S2.**
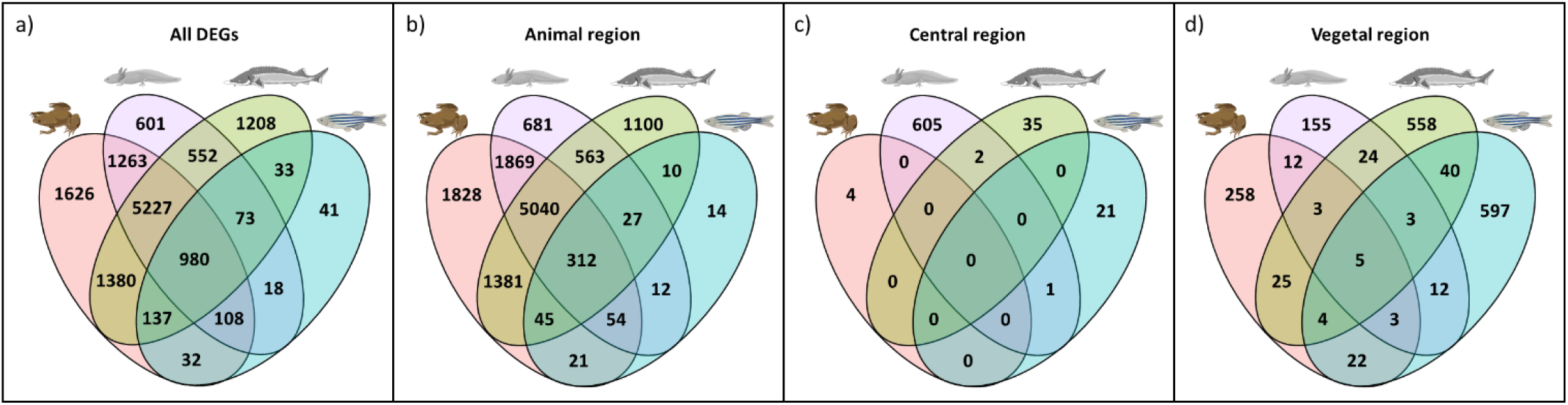
Overlap of unique maternal DEGs (dataset1) with annotated gene symbols between the different models. a) all maternal DEGs, b) extreme animal and animal localized DEGs, c) centrally localized DEGs, d) extreme vegetal and vegetal localized DEGs.

**Supplementary file 1: Figure 3.**
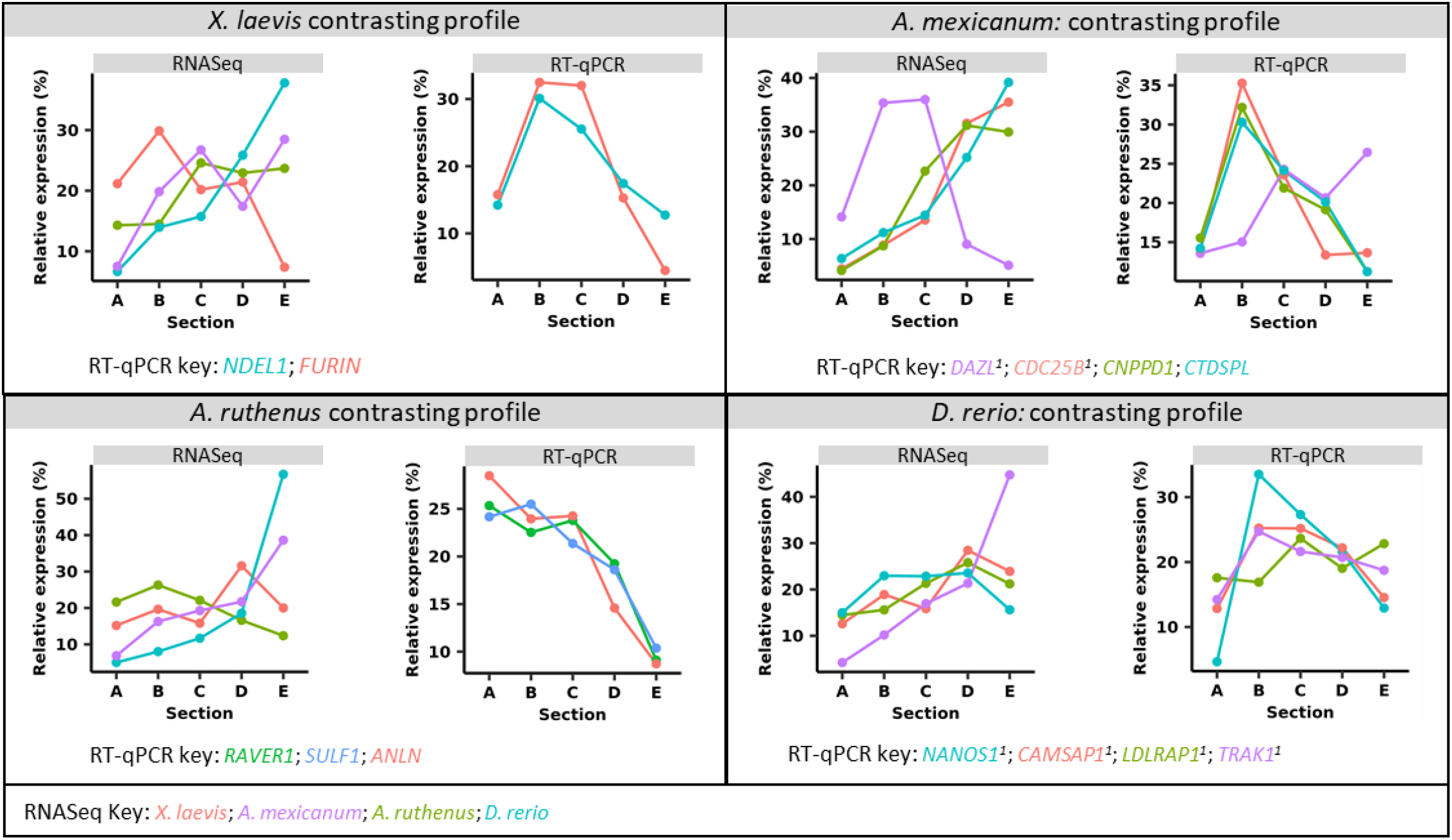
RNA-Seq expression profiles of genes that were found to be exclusively vegetally localized in three models but an alternative location in the fourth model. The RNA-Seq profiles represent the medium expression for the given set of genes per model. The RT-qPCR shows the averaged expression profiles for each of the contrasting genes for the given model. The RT-qPCR profiles for many of the contrasting genes were either animal, animal-central or showed high variability between replicates. ^1^ – high variability between replicates

**Supplementary file 1: Figure S4.**
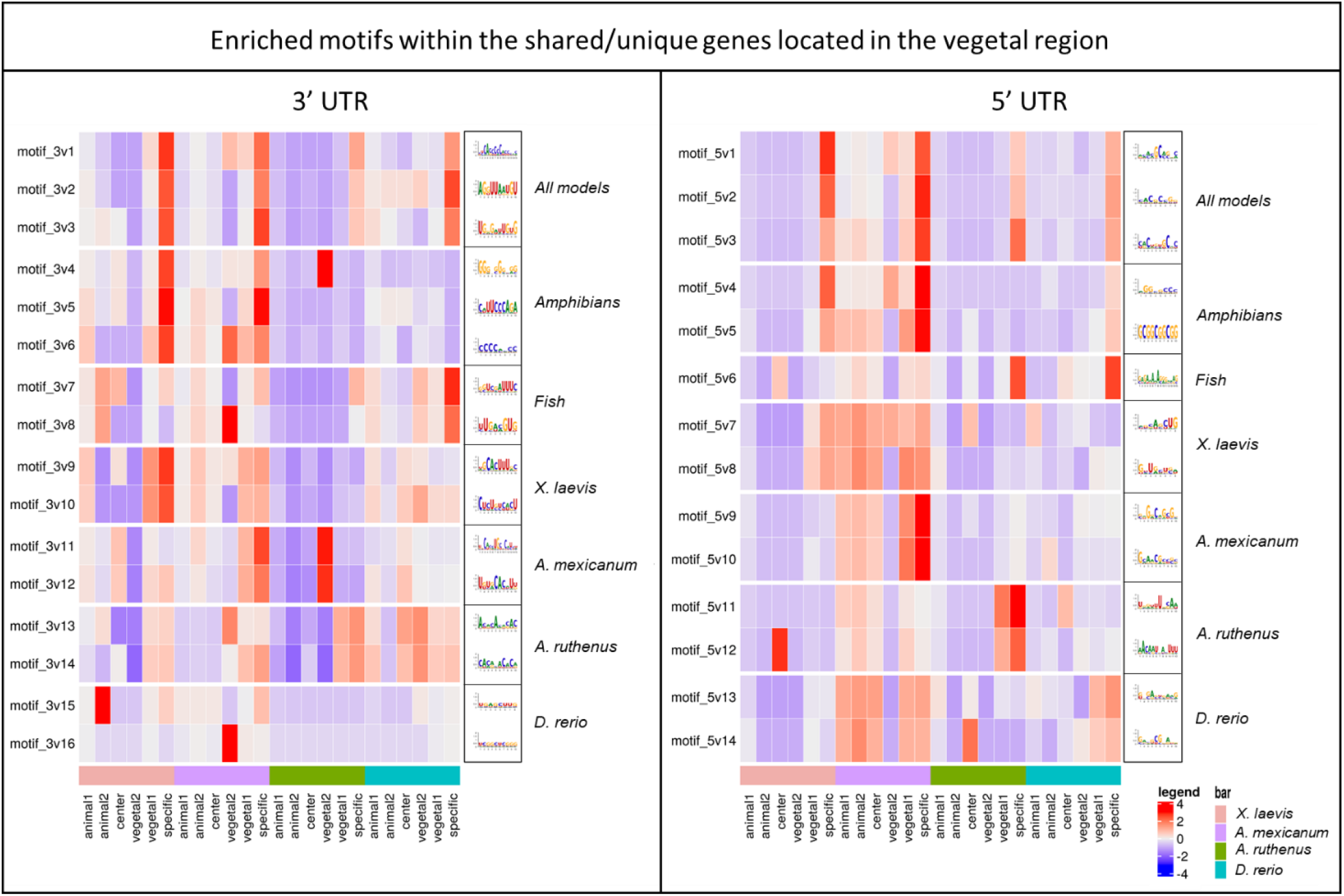
Heatmap of the z-score showing the proportion of genes from each localization profile that contained the given vegetal motif as derived using FIMO. The shown motifs represent those that are statistically significantly enriched within the genes of interest (specific) versus the contrasting profiles from dataset2 (animal1, animal2, central) and also 2x more abundant versus these same contrasting categories. Member genes within the “specific” category can be found in the dataset3. The gene representatives for the given localization categories represent those from the smaller pool of DEGs that showed the best reproducibility (dataset2). Animal1 contains extreme animal and animal profiles that showed peak in either section A, A and B or B of the egg; animal2 contains animal profiles that showed peak in section B and C; center contains profiles that showed peak in section C; vegetal2 contains vegetal profiles that showed peak in section C and D; vegetal1 contains extreme vegetal and vegetal profiles that showed peak in either section D, D and E, or E of the egg.

**Supplementary file 1: Figure S5.**
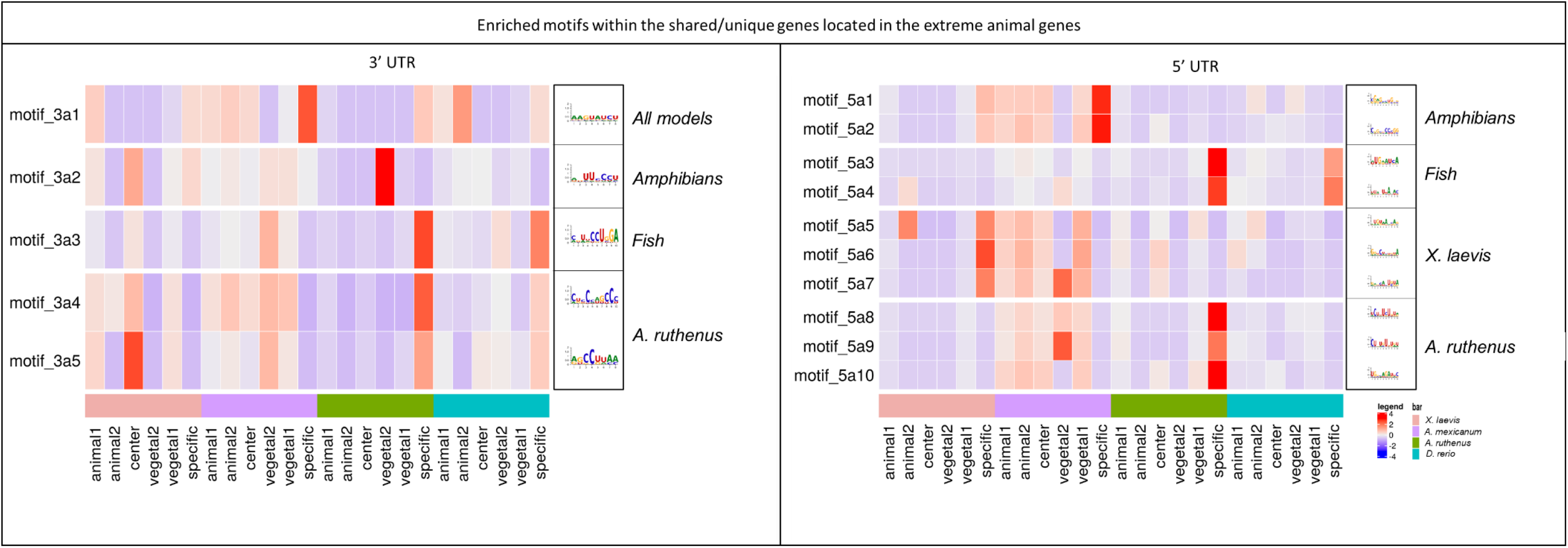
Heatmap of the z-score showing the proportion of genes from each localization profile that contained the given extreme animal motif as derived using FIMO. The shown motifs represent those that are statistically significantly enriched within the genes of interest (specific) versus the contrasting profiles from dataset2 (animal1, animal2, central) and also 2x more abundant versus these same contrasting categories. Member genes within the “specific” category can be found in the dataset3. The gene representatives for the given localization categories represent those from the smaller pool of DEGs that showed the best reproducibility (dataset2). Animal1 contains extreme animal and animal profiles that showed peak in either section A, A and B or B of the egg; animal2 contains animal profiles that showed peak in section B and C; center contains profiles that showed peak in section C; vegetal2 contains vegetal profiles that showed peak in section C and D; vegetal1 contains extreme vegetal and vegetal profiles that showed peak in either section D, D and E, or E of the egg.

**Supplementary file 1: Figure S6.**
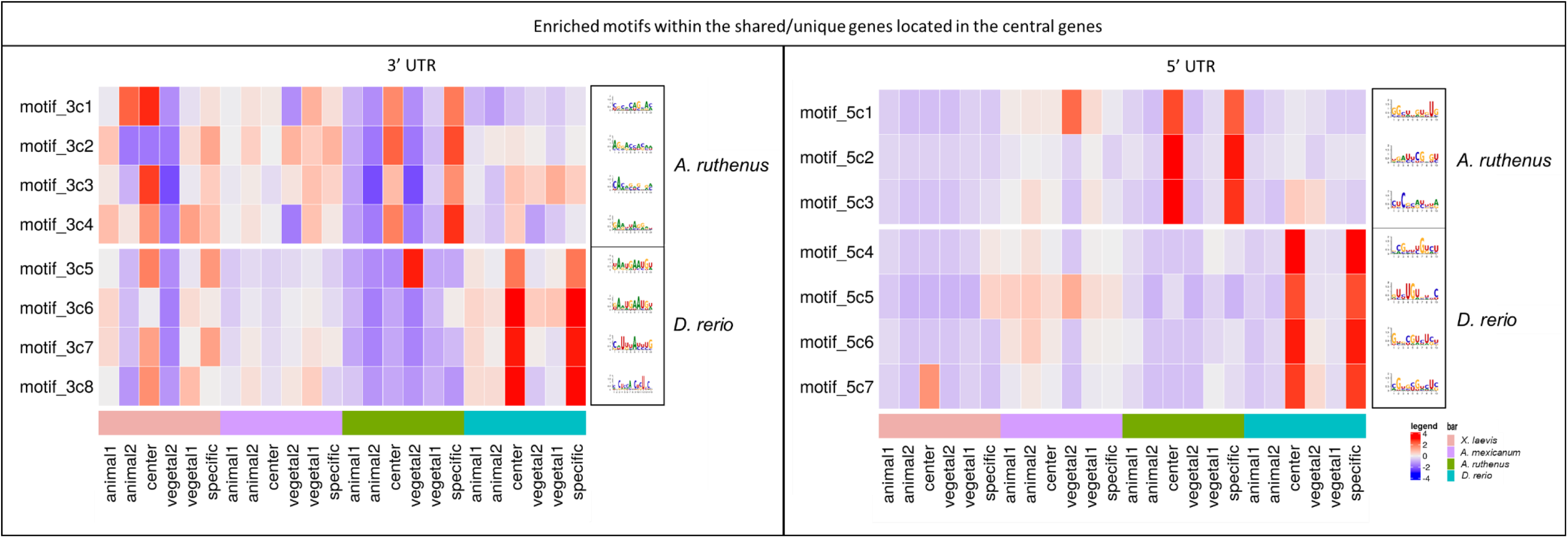
Heatmap of the z-score showing the proportion of genes from each localization profile that contained the given central motif as derived using FIMO. The shown motifs represent those that are statistically significantly enriched within the genes of interest (specific) versus the contrasting profiles from dataset2 (animal1, animal2, central) and also 2x more abundant versus these same contrasting categories. Member genes within the “specific” category can be found in the dataset3. The gene representatives for the given localization categories represent those from the smaller pool of DEGs that showed the best reproducibility (dataset2). Animal1 contains extreme animal and animal profiles that showed peak in either section A, A and B, or B of the egg; animal2 contains animal profiles that showed peak in section B and C; center contains profiles that showed peak in section C; vegetal2 contains vegetal profiles that showed peak in section C and D; vegetal1 contains extreme vegetal and vegetal profiles that showed peak in either section D, D and E, or E of the egg.

**Supplementary file 1: Figure S7.**
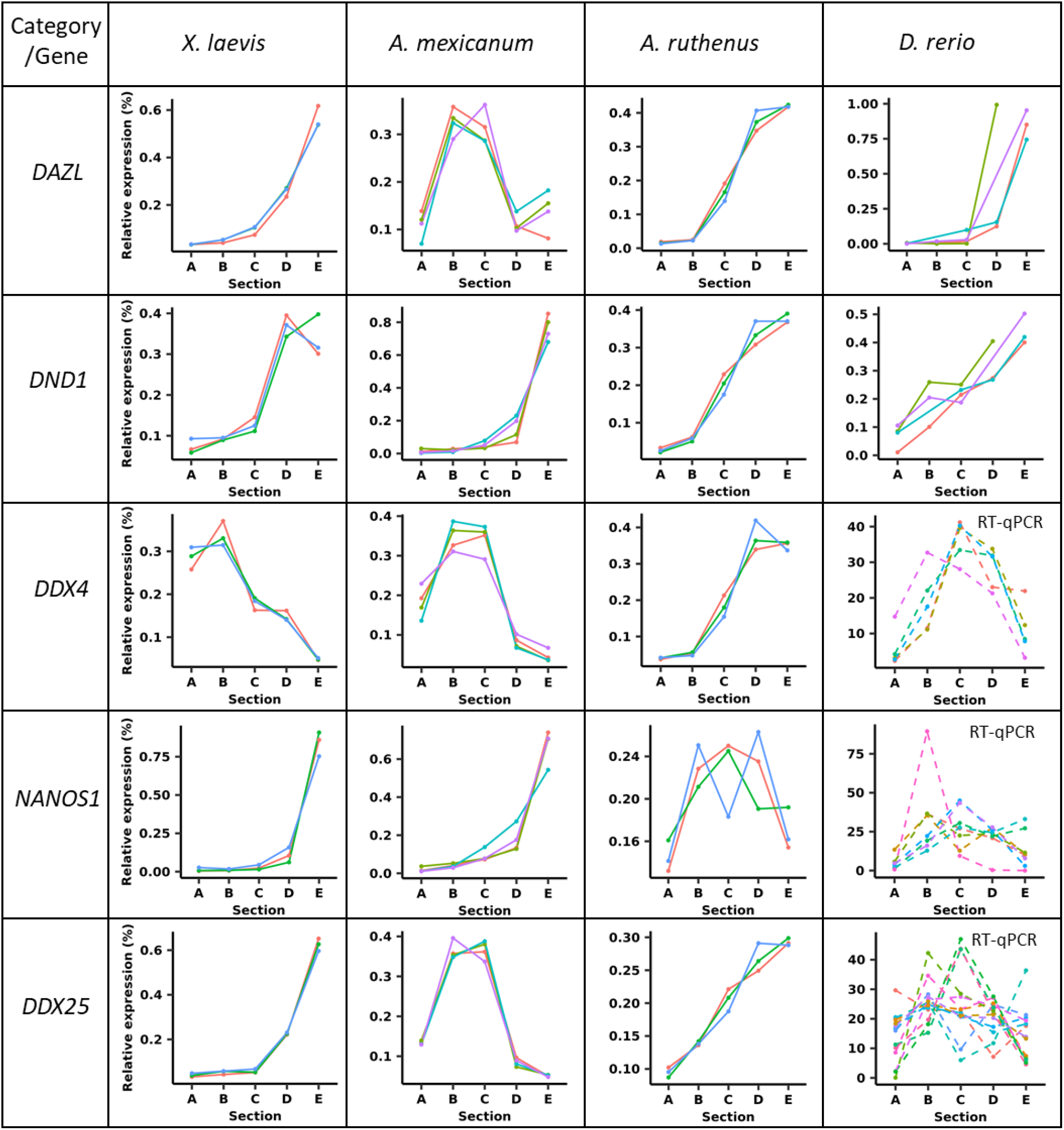
Localization profiles for some key member genes belonging to PGC markers. Graphs with dashed lines indicate that the RNASeq data for the gene was not differentially expressed, failed the statistical analysis using DESeq or did not meet threshold criteria. Graphs marked with RT-qPCR represent replicate data from RT-qPCR assays, while all other unmarked graphs show replicate data from the TOMO-Seq.

**Supplementary file 1: Figure S8.**
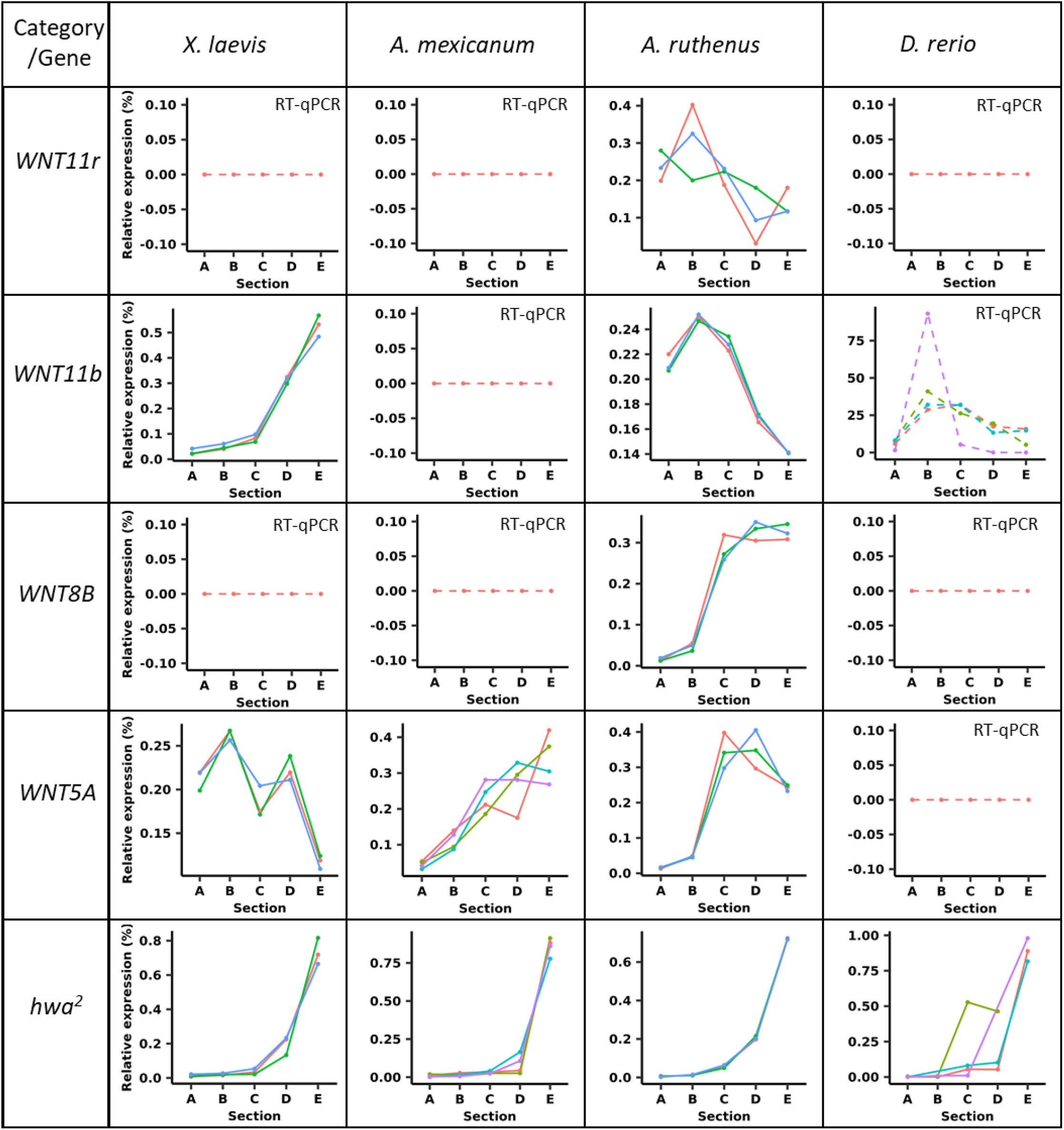
Localization profiles for some key member genes belonging to wnt ligands. Graphs with dashed lines indicate that the RNA-Seq data for the gene was not differentially expressed, failed the statistical analysis using DESeq or did not meet threshold criteria. Graphs marked with RT-qPCR represent replicate data from RT-qPCR assays, while all other unmarked graphs show replicate data from the TOMO-Seq. ^2^gene nomenclature for *D. rerio*.

**Supplementary file 1: Figure S9.**
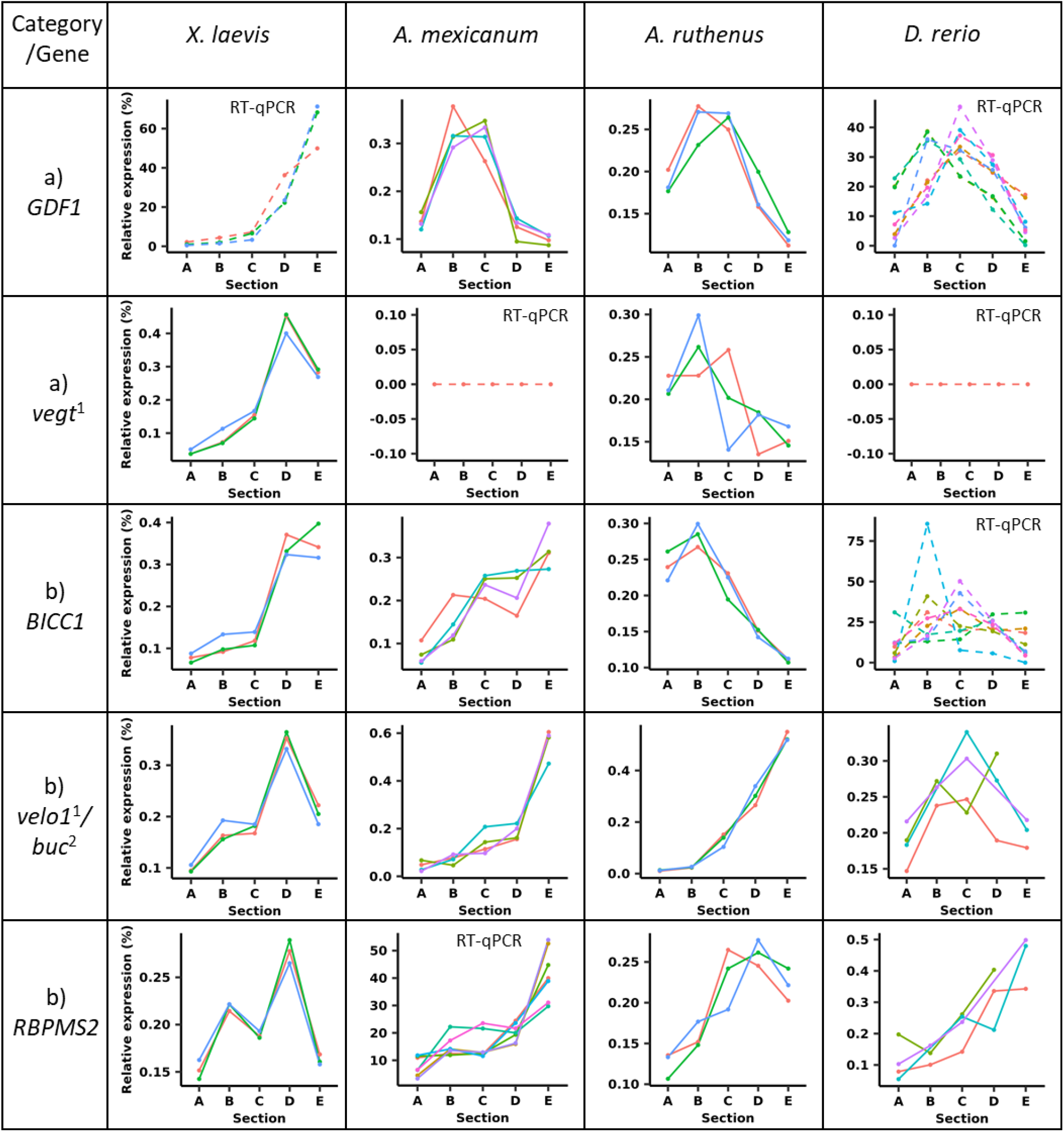
Localization profiles for some key member genes belonging to a) Endodermal and mesodermal determinants, and other b) known *Xenopus laevis* vegetal genes. Graphs with dashed lines indicate that the RNA-Seq data for the gene was not differentially expressed, failed the statistical analysis using DESeq or did not meet threshold criteria. Graphs marked with RT-qPCR represent replicate data from RT-qPCR assays, while all other unmarked graphs show replicate data from the TOMO-Seq. *Danio rerio GDF3* gene is an ortholog of *GDF1* gene. *velo^1^* (amphibians and *A. ruthenus*) and *buc^2^* (*D. rerio*) are orthologous genes. ^1^gene nomenclature for *X. laevis;* ^2^gene nomenclature for *D. rerio*.

**Supplementary file 1: Figure S10.**
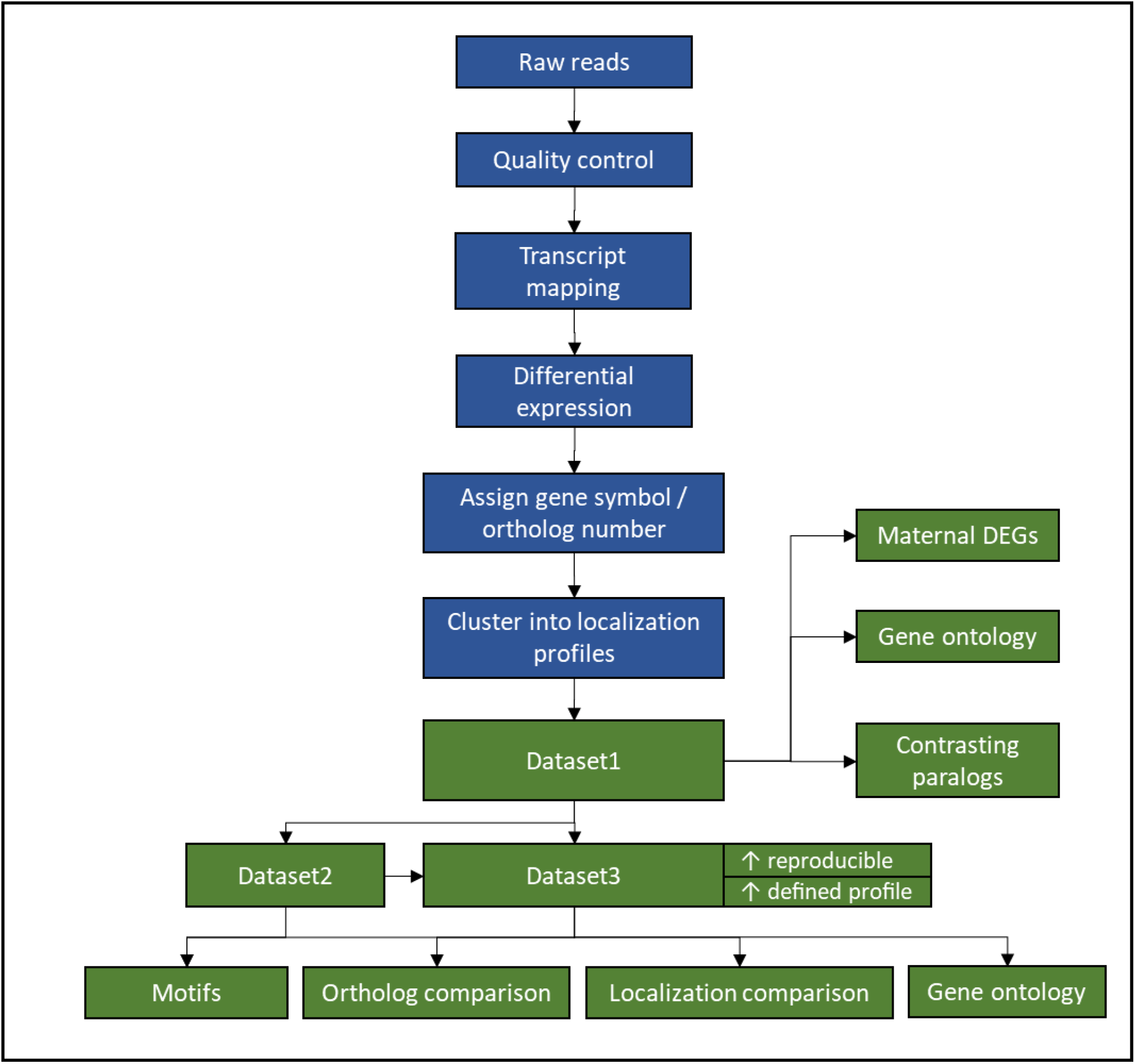
Summary flowchart showing the RNASeq data analysis process.

### Supplementary file 2

***Supplementary file 2: Table S1.* Datasets showing differentially expressed genes (DEGs) for each model.** The symbols are derived from the most likely human ortholog symbol or in some instances a given model’s symbol. Dataset1 shows all DEGs. Dataset2 represents reproducible DEGs with well-defined profiles and was used for the detection of known motifs and also motif enrichment analysis. Dataset3 also comprises DEGs with well-defined profiles but also those whose annotation/orthology was further analysed in detail. The dataset3 was used for more detailed ortholog and localization comparative analysis between the models and gene ontology analysis.

***Supplementary file 2: Table S2.* List of differentially expressed genes whose paralogs show contrasting localization profiles (one form has extreme vegetal/vegetal profile and the duplicated form has extreme animal/animal profile).**

***Supplementary file 2: Table S3.* Vegetal (extreme vegetal & vegetal) differentially expressed genes (only those with symbols) - dataset3, that are shared across all models, among three model, the amphibians, the fishes or unique to each model.**

***Supplementary file 2: Table S4.* Enrichment analysis of Gene Ontologies, biological pathways, regulatory motifs and protein complexes associated with conserved extreme vegetal and vegetal genes (dataset3).**

***Supplementary file 2: Table S5.* Central differentially expressed genes (only those with symbols) - dataset3, that are shared among the models or unique to each model.**

***Supplementary file 2: Table S6. Enrichment analysis of Gene Ontologies, biological pathways, regulatory motifs and protein complexes associated with conserved central genes (dataset3).***

***Supplementary file 2: Table S7.* Extreme Animal differentially expressed genes (only those with symbols) - dataset3, that are shared across all models, the amphibians, the fishes or unique to each model.**

***Supplementary file 2: Table S8.* Enrichment analysis of Gene Ontologies, biological pathways, regulatory motifs and protein complexes associated with conserved extreme animal genes (dataset3).**

***Supplementary file 2: Table S9.* Vegetal motifs that are at least 2x abundant in the extreme vegetal/vegetal genes of interest and also statistically significantly enriched relative to one of the other sections.**

***Supplementary file 2: Table S10.* Central motifs that are at least 2x abundant in the central genes of interest and also statistically significantly enriched relative to one of the other sections.**

***Supplementary file 2: Table S11.* Extreme animal motifs that are at least 2x abundant in the extreme animal genes of interest and also statistically significantly enriched relative to one of the other sections.**

***Supplementary file 2: Table S12.* Published localization profiles of certain genes based on *in situ*.**

***Supplementary file 2: Table S13.* Primer pairs used for the verification of the localization profiles of selected maternal genes.**

***Supplementary file 2: Table S14.* Summary of the quantity of RNA, kits used during library preparation and basic setup of the RNA-Seq sequencing used for each model.**

***Supplementary file 2: Table S15.* Summary of some basic quality control and alignment parameters for the RNA-Seq fragments.**

***Supplementary file 2: Table S16.* Marker genes used for NormQ.**

***Supplementary file 2: Table S17.* Algorithm for localization profiles for dataset1.**

***Supplementary file 2: Table S18.* Algorithm criteria to define dataset2 for use in motif detection and motif enrichment analysis.**

